# Artemis Regulates Homology-Independent Prime Editing (PRINS) for Enhanced Genomic Insertions

**DOI:** 10.64898/2025.12.08.693005

**Authors:** Louis C. Dacquay, Oi Kuan Choong, Angela Membrino, Anders Lundin, Nina Akrap, Johannes Köllner, Pei-Pei Hsieh, Anna Baran, Euan Gordon, Salman Mustfa, Sabina Saka, Danang Crysnanto, Fredrik Karlsson, Anders Dahlen, George Thom, Julia Lindgren, Mike Firth, Marie K. Schwinn, Thomas Machleidt, Martin Peterka, Sasa Svikovic, Grzegorz Sienski, Marcello Maresca

**Author notes:** Correspondence to: Louis C. Dacquay, Marcello Maresca.

## Abstract

Nuclease-based prime editing (PEn) offers enhanced genomic insertion efficiency compared to nickase-based prime editors, but its reliance on double-strand break (DSB) repair leads to complex and often unpredictable on-target indel distributions. PRINS editing, a PEn variant utilizing springRNAs, uniquely relies on non-homologous end joining (NHEJ) for insertions, providing an insertion-only strategy ideal for functional protein tagging or serine integrase landing pad insertions, yet it suffers from inherent imprecision. Here, we identify the DNA repair factor Artemis (*DCLRE1C*) as a key regulator of PEn-generated indel profiles, particularly in springRNA-mediated PRINS editing.

Through a targeted genetic screen, we show that the absence of Artemis significantly shifts indel distributions away from deletions and shorter truncations towards longer, functionally acceptable insertions. Our data indicates that Artemis cleaves PEn-generated 3’-overhangs in a length-dependent manner, with its impact increasing for longer reverse-transcribed overhangs. This understanding reveals that regulating Artemis activity can improve insertion frequency specifically for PRINS. We develop and validate robust epigenetic (CRISPRoff) and antisense oligonucleotide (ASO) strategies to effectively silence/knockdown Artemis expression, successfully recapitulating the beneficial PRINS editing outcomes observed in Artemis-deficient cells. Leveraging these insights, we show that Artemis modulation can enhance endogenous protein tagging in cells, including challenging hiPSC-derived non-dividing cardiomyocytes. Our findings support Artemis as a key regulator of PRINS editing outcomes and present a tunable strategy to optimize insertion efficiency for diverse genomic engineering and therapeutic applications.

## Introduction

The ability to edit the DNA sequence of the genome with CRISPR-Cas9 systems has revolutionized biological research and holds immense promise for therapeutic or translational applications. Prime editing (PE) has emerged as a significant technological advancement, enabling versatile “search-and-replace” gene editing strategy for diverse types of edits into precise regions of the genome [1]. This technology relies on the fusion of a Cas9 nickase, with its HNH nuclease domain deactivated, with a reverse transcriptase (RT), used in conjunction with a modified guide RNA (pegRNA) that includes the spacer and the gRNA scaffold for Cas9 recognition and genome targeting, as well as a 3’ extension that acts as the template for the reverse transcriptase.

More recently, nuclease-based prime editing (PEn) has further expanded the capabilities of prime editing [2–5]. PEn utilizes a fully active Cas9 nuclease to generate a DSB at the target site alongside a pegRNA, a strategy that can improve prime editing efficiency, particularly at sites that are poorly edited by conventional nickase-based prime editors [5]. Apart from human cell editing, PEn editing strategies have been shown to be quite versatile in different applications. For example, inserting nuclear localization signal sequences in zebrafish [6], inserting protein epitope tags in rice [7], and as a selective enrichment system for various modifications in bacteria [8]. Within the PEn framework, a distinct strategy known as PRINS editing (PRImed INSertion strategy) employs a specialized springRNA that, unlike the pegRNA, omits the homology arm [5]. This design directs the 3’-overhang generated by PEn to primarily rely on the host cell’s non-homologous end joining (NHEJ) pathway for insertion. PRINS editing offers unique advantages: as an insertion-only strategy, it is well-suited for applications that only require the introduction of new genetic material and is particularly beneficial in cells that are mostly reliant on NHEJ activity to repair DSBs, such as homology directed repair (HDR)-deficient or non-dividing cells. This makes it a useful tool for applications in these types of cells especially in-frame insertion of protein epitope tags (*e.g.*, FLAG, HIBIT) or the integration of serine integrase attachment sites (*e.g.*, attB sequences for Bxb1 integrase), facilitating downstream genetic manipulation with large DNA cargoes.

Despite these promising applications, PRINS editing also faces significant drawbacks stemming from its reliance on the inherently imprecise NHEJ pathway. The non-homologous end joining (NHEJ) pathway is the primary mechanism for repairing DSBs in mammalian cells, directly ligating broken DNA ends, often with minimal or no sequence homology. This process involves a hierarchical assembly of protein factors. The Ku heterodimer (Ku70/Ku80) initially binds to DSB ends, recruiting the catalytic subunit of DNA-dependent protein kinase (DNA-PKcs, encoded by *PRKDC*). DNA-PKcs then phosphorylates various substrates, including itself and Artemis. Artemis (*DCLRE1C*), a nuclease whose activity is stimulated by DNA-PKcs, is crucial for processing complex DNA end structures, such as hairpin loops or incompatible overhangs, to generate ligation compatible ends together with the polymerase activity of Polλ and Polµ. Following end processing, the Xrcc4-DNA ligase IV-XLF (*NHEJ1*) complex catalyzes the final ligation step. Additionally, 53BP1 (*TP53BP1*) functions upstream of this pathway by recognizing DSBs and promoting NHEJ by antagonizing homologous recombination [9]. A major challenge for PEn is the lack of control over editing outcomes, as the 3’-overhang processing by NHEJ can result in a wide spectrum of indels beyond the desired insertion.

These include deletions, truncated insertions, and incorporation of the gRNA scaffold, making it difficult to achieve desired insertion events. The complexity of these indel distributions necessitates a deeper understanding of the host cellular DNA repair machinery involved.

Here, we report that the DNA repair factor Artemis (*DCLRE1C*) is a critical regulator of PEn-generated indel profiles, particularly for springRNA-mediated PRINS editing. Through a targeted genetic screen, we identified Artemis as a key determinant of insertion outcomes, demonstrating that its absence significantly shifts indel distributions away from deletions and truncated products towards longer, more desired insertions, especially for PRINS editing. We uncover that Artemis cleaves PEn-generated 3’-overhangs in a length-dependent manner, demonstrating that its regulation can be leveraged to improve insertion frequency. Furthermore, we develop and validate robust epigenetic and antisense oligonucleotides (ASOs) strategies to achieve effective Artemis silencing or knockdown. We demonstrate that Artemis knockdown enhances the efficiency of PRINS editing for functional tagging of endogenous genes including in hiPSC-derived non-dividing cardiomyocytes. Our findings thus establish a tunable strategy to optimize PEn-mediated insertion outcomes for diverse genomic engineering and therapeutic applications.

## Results

### Targeted Genetic Screen Identifies Artemis and NHEJ Factors as Key Modulators of PEn-generated Indel Profiles

Nuclease-based prime editing (PEn) enhances genomic insertion efficiency but generates a complex on-target indel landscape. To optimize desired insertions, especially for homology-independent PRINS editing, understanding host DNA repair is critical. Building on previous genetic screens identifying mismatch repair (MMR) genes as modulators of nicking-based prime editing [10, 11], we performed a targeted screen of 33 DNA repair gene knockouts in leukemia-derived HAP1 cells ([12], Horizon Discovery). Our objective was to identify factors modulating insertion frequencies and their precise nature for both pegRNA-mediated PEn and springRNA-mediated PRINS editing, assessing relative editing outcomes including non-templated insertions and deletions (indels), overall prime editing events, and precise 6 base pairs (bp) templated insertion at the AAVS1 (*PPP1R12C*) target site. In contrast to findings for nicking-based PE which were affected by mismatch repair genes, our screen revealed that NHEJ pathway factors—*LIG4*, *NHEJ1*, *XRCC4*, *PRKDC*, *TP53BP1*, and *DCLRE1C* (Artemis)—exerted the most prominent effects (Figure 1a). Allele frequency plots (Supplementary Figure 1) confirmed that these insertion events were pegRNA/springRNA templated, and not random insertions or genome duplication events. To further characterize these effects and showcase how different NHEJ components distinctly modulate PEn outcomes, we analyzed representative knockout lines (WT, *LIG4*, *TP53BP1*, and *DCLRE1C*) through indel distribution plots (Figure 1b).

**Figure 1.**
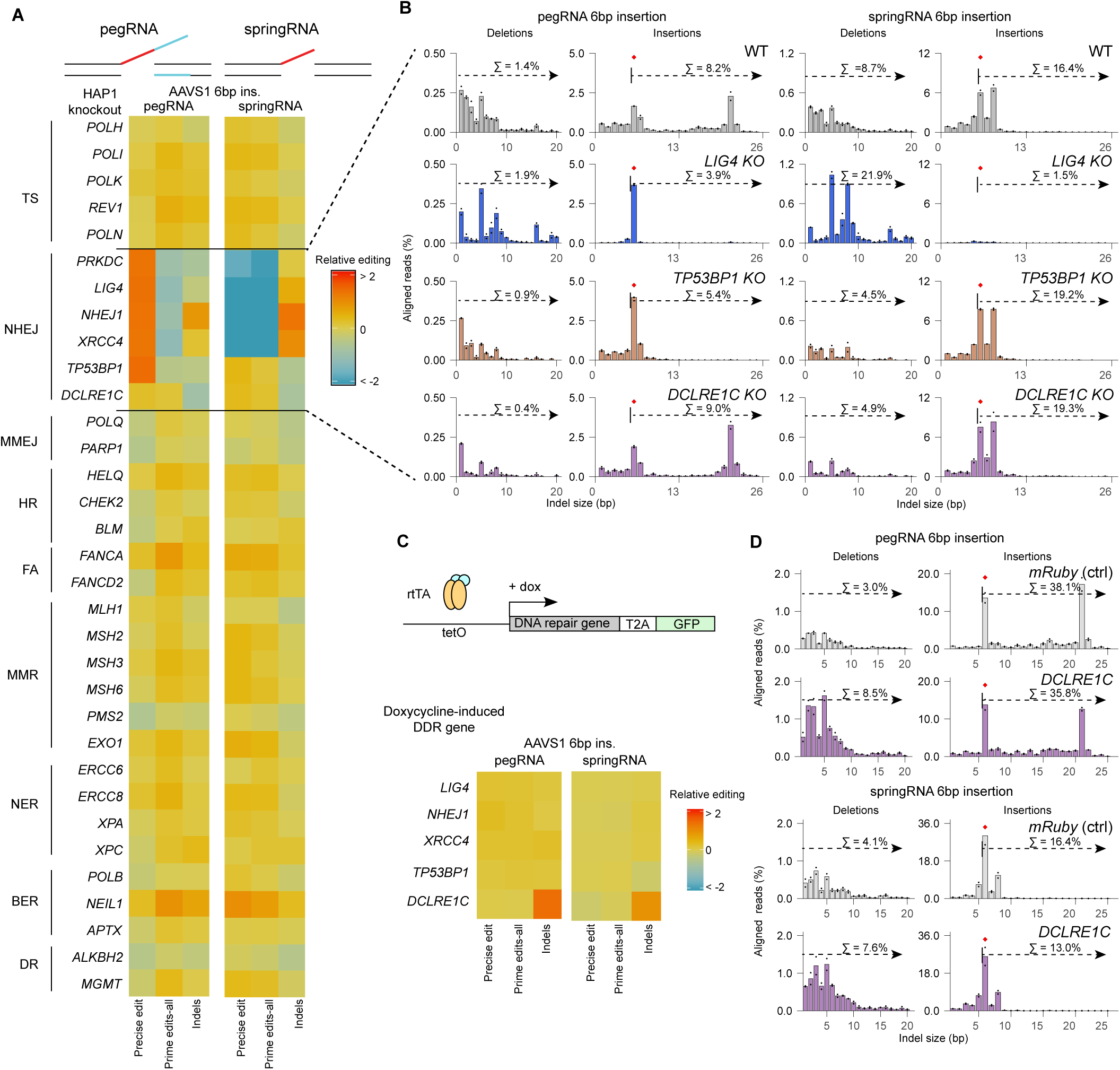
Targeted Genetic Screen Identifies Artemis and NHEJ Factors as Key Modulators of PEn-generated Indel Profiles. A) HAP1 knockout library screen of 33 DNA repair factor lines and wild-type (WT) cells. Cells were electroporated with *Sp*Cas9-RT (PEn) RNPs and either pegRNA (with homology arm) or springRNA (without homology arm) designed for a 6 bp insertion at the AAVS1 (*PPP1R12C*) target site. Editing outcomes were analyzed by amplicon-seq and quantified using CRISPResso2 using‘Prime editing’ mode.’Precise edits’ are unmodified prime edits;’Prime edits-all’ includes all prime edits and scaffold incorporations;’Indels’ are modified reference reads. Editing percentages for these categories were normalized to total editing efficiency, then made relative to HAP1 WT, and log2-transformed. Heatmap displays average relative frequencies (n=2 biological replicates), grouped by DNA repair pathway, with red indicating higher and blue lower frequency (saturated at ± 2 log2 fold change). TS = Translesion Synthesis, NHEJ= Non-Homologous End Joining, MMEJ= Microhomology Mediated End Joining, HR = Homologous Recombination, FA = Fanconi Anemia, MMR = MisMatch Repair, NER = Nucleotide Excision Repair, BER = Base Excision Repair, DR = Direct Reversal B) Indel distribution patterns for PEn editing (with pegRNA or springRNA) in WT and representative NHEJ knockout HAP1 cells (*DCLRE1C*, *TP53BP1*, *LIG4*). Indel histograms, generated by CRISPResso2 and a custom R-script, show average indel size frequencies (n=2 biological replicates). Deletions (1-20 bp) are plotted on the left, and insertions (1-26 bp) on the right. Intended insertion size is indicated by a red diamond. Editing frequencies for all deletion events and for all intended or longer insertion events are calculated by taking the sum of the indicated editing events from the individual indel size frequencies from the analysis described above (Σ deletions = indel size-1 to-min, Σ intended or longer insertions = indel size 6 to max). The numerical value on each graph is the average of the two biological replicates. C) Effect of NHEJ gene overexpression on PEn editing outcomes. HEK293T cell lines were generated with Xential (see methods) with an integrated doxycycline-inducible system to overexpress either exogenous mRuby (control) or individual NHEJ genes (*LIG4*, *NHEJ1*, *XRCC4*, *DCLRE1C*, *TP53BP1*). Cells were first treated with doxycycline (2 µg/mL) for one day to induce expression of cDNA, then electroporated with *Sp*Cas9-RT (PEn) RNPs and either pegRNA or springRNA designed for a 6 bp insertion at the AAVS1 (*PPP1R12C*) target site. Editing analysis was performed as described in (A). The heatmap displays average relative frequencies (n=2 biological replicates) for the five NHEJ overexpressing cell lines compared to the mRuby control, using the same color scale and saturation as in (A). RT= M-MLV reverse transcriptase; DDR = DNA Damage Repair; rtTA = reverse tetracycline-controlled transcriptional activator; tetO = tetracycline operons; GFP = Green Fluorescent Protein D) Indel distribution patterns for PEn editing comparing mRuby (control) and *DCLRE1C* overexpressed HEK293T backgrounds. Indel distribution plots were generated as described in (B). Intended insertion size is indicated by a red diamond.

Indel distributions across knockout lines highlighted distinct roles for NHEJ components. Knockout of *LIG4*, a core NHEJ factor along with *XRCC4* and *NHEJ1*, generally decreased NHEJ-mediated insertions relative to wild-type, while increasing frequency of homology-dependent, precise 6 bp pegRNA insertions. For springRNA editing, intended or longer insertions were substantially reduced in the *LIG4* knockout, decreasing from 16.4% in wild type to 1.5%. Deletion events increased, particularly for springRNA (from 8.7% to 21.9%), and the deletion size distribution shifted, consistent with a greater contribution from microhomology-mediated end joining (MMEJ). Similar to *LIG4* knockout, *TP53BP1* knockout increased pegRNA-mediated precise insertions at the expense of longer unintended insertions. However, unlike *LIG4* knockout, *TP53BP1* knockout slightly increased springRNA insertion frequencies, raising intended or longer insertions from 16.4% to 19.2%, suggesting that *TP53BP1* promotes homology-driven repair of the pegRNA homology arm rather than broadly inhibiting NHEJ. *TP53BP1* knockout also decreased deletions for both pegRNA (from 1.4% to 0.9%) and springRNA (from 8.7% to 4.5%) editing.

Artemis knockout (*DCLRE1C*) consistently decreased deletions compared to wild-type for both pegRNA (from 1.4% to 0.4%) and springRNA (from 8.7% to 4.9%) editing. We also observed a subtle shift in insertion profiles away from smaller truncated products toward intended or longer insertions, increasing their cumulative frequency from 8.2% to 9.0% (pegRNA) and from 16.4% to 19.3% (springRNA). While this suggests a role for Artemis in shaping insertion length, the effect is modest at for this 6 bp insertion target.

To confirm direct genetic involvement, we examined overexpression of NHEJ-factors in HEK293T cells with a doxycycline-inducible system integrated into the *HBEGF* locus [13] (Figure 1c). Only exogenous overexpression of *DCLRE1C* influenced PEn outcomes compared to the mRuby control conditions while overexpression of *TP53BP1* and other factors did not produce substantial effects. Overexpressing Artemis had the opposite effect as the knockout, increasing deletion events from 3.0% to 8.5% (pegRNA) and from 4.1% to 7.6% (springRNA) (Figure 1d). It also subtly decreased the cumulative frequency of intended or longer insertions from 38.1% to 35.8% (pegRNA) and from 16.4% to 13.0% (springRNA).

As a control to test whether the overexpression phenotype requires Artemis nuclease activity, we overexpressed catalytically dead variants of Artemis and assessed springRNA-mediated PRINS editing. Although catalytically dead Artemis mutants have been reported to act as dominant negatives in cancer cells (increasing radiosensitivity characteristic of Artemis deficiency) [14, 15], in our system overexpression of catalytically inactive Artemis did not alter editing relative to the mRuby control (Supplementary Figure 2). By contrast, overexpression of wild-type *DCLRE1C* produced the expected decrease in intended or longer insertions and increase in deletions. These results demonstrate that the overexpression effect depends on a functional nuclease domain, and that catalytically inactive Artemis does not exert dominant negative effects on PRINS editing under our conditions.

These initial findings support Artemis as a regulator of PEn outcomes, particularly in balancing deletion and insertion length, providing a motivation for further investigation.

### Artemis Deficiency Promotes Longer Insertions with homology-independent PRINS Editing

The initial 6 bp insertion screen hinted at Artemis’s influence on insertion length, although this effect did not appear that pronounced at first glance. To better characterize and generalize this, we examined Artemis (*DCLRE1C*) knockout in HAP1 cells at four additional genomic sites (*DPM2*, *PCSK9*, *PDCD1*, *TRAC*) for longer desired insertions (8-12 bp), assessing both pegRNA and springRNA editing.

Remarkably, *DCLRE1C* knockout consistently shifted the indel distribution of homology-independent springRNA editing towards longer insertions, reducing both deletions and shorter truncated insertion products across all four target sites (Figure 2a, Supplementary Figure 3). Homology-dependent pegRNA editing showed some sensitivity to *DCLRE1C* knockout, but competing homology-driven repair from the homology arm likely buffered the effect of NHEJ-mediated 3’-overhang processing. The effect of Artemis knockout on the relative insertion and deletion frequency was also recently observed in a genome wide profiling of gene knockouts on Cas9-induced DSB repair [16], but this effect seems more pronounced here with the longer 3’-overhang generated by PRINS than with the blunt or short overhang ends generated by *Sp*Cas9.

**Figure 2.**
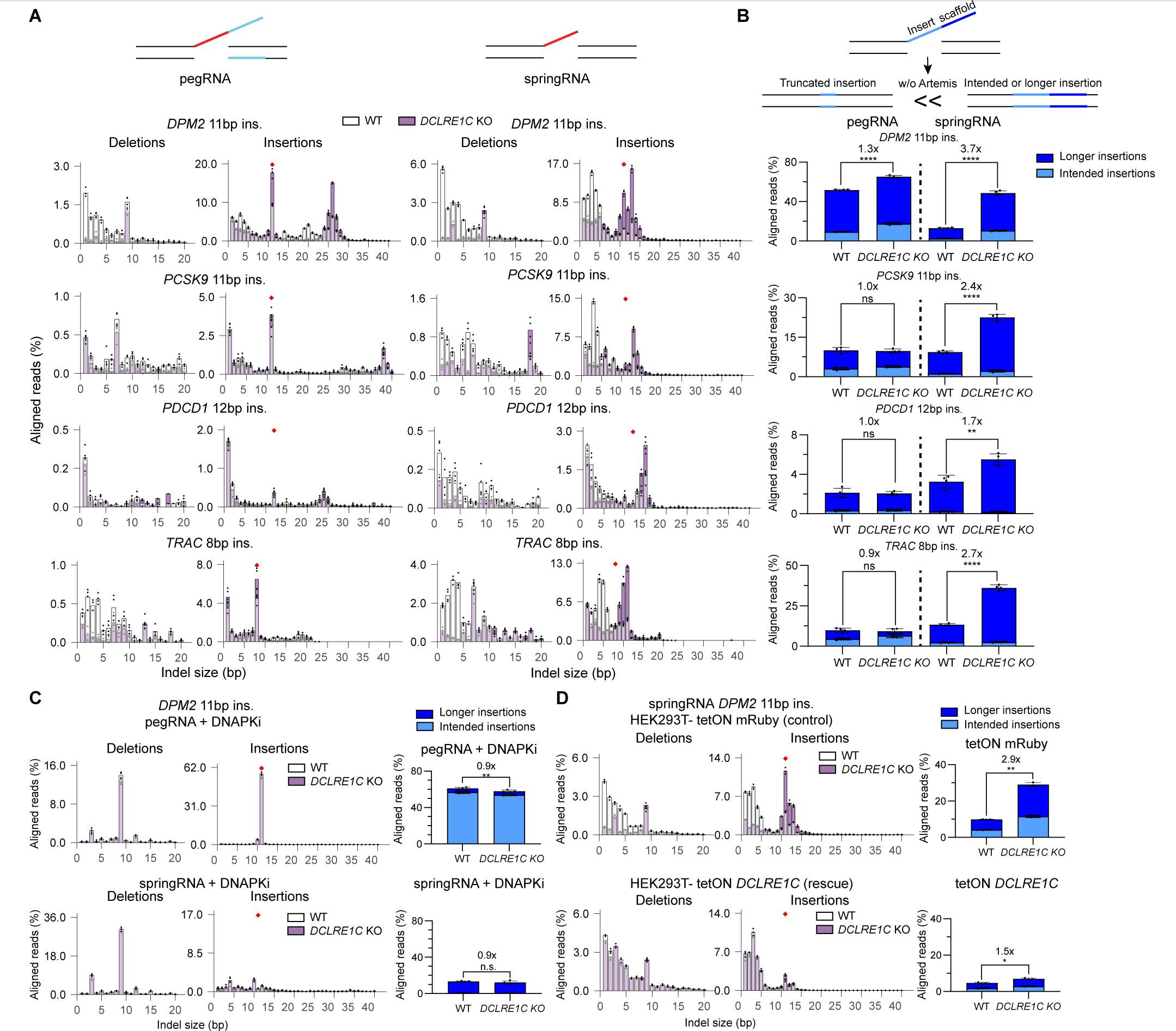
Artemis Deficiency Promotes Longer Insertions with Homology-Independent PRINS Editing. A) Indel distribution patterns for PEn editing with pegRNA (left panel) or springRNA (right panel) for longer insertions in wild-type (WT) and *DCLRE1C* knockout HAP1 cells. Cells were electroporated with *Sp*Cas9-RT (PEn) RNPs and either pegRNA or springRNA for each target site. Indel histograms, generated by CRISPResso2 and a custom R-script, show average indel size frequencies (n=4 biological replicates) between WT (white) and *DCLRE1C* knockout (purple) overlayed on top of each other, at four target sites (*DPM2*-11bp, *PCSK9*-11bp, *PDCD1*-12bp, and *TRAC*-8bp) with indicated insertion size. Deletions (1-20 bp) are plotted on the left, and insertions (1-40 bp) on the right. Intended insertion size is indicated by a red diamond. B) Insertion frequencies of PEn editing between WT and *DCLRE1C* knockout HAP1 cells. Schematic (top) illustrates effect on Artemis on insertion types. Presence of Artemis leads to truncated insertions, while its absence leads to increased intended or longer insertions. Bar graph shows average frequency of intended or longer insertions (four biological replicates ± SD) from the above experiment, calculated using CRISPResso2 and a custom R-script. C) DNA-PKcs inhibitor (DNAPKi) inhibits the effect of *DCLRE1C* knockout HAP1 cells. Cells were electroporated with *Sp*Cas9-RT (PEn) RNPs and either pegRNA or springRNA for the indicated target site. Electroporated cells were directly seeded in plates containing DNAPKi (1 µM) or DMSO as a control. Overlayed indel distribution patterns (left) and insertion frequencies (right) show results of PEn editing (with pegRNA or springRNA) at the *DPM2* target site (11 bp intended insertion) in WT and *DCLRE1C* knockout HAP1 cells, in the presence of DNAPKi. Data analyzed as described in (A) and (B) (n=4 biological replicates ± SD). Intended insertion size on histograms is indicated by a red diamond. D) Overexpression of exogenous *DCLRE1C* reverts the effect of *DCLRE1C* knockout. HEK293T cell lines (WT and *DCLRE1C* knockout) were generated by Xential (see methods) with a doxycycline-inducible system to overexpress either mRuby (control) or *DCLRE1C* cDNA. Cells were first treated with doxycycline (2 µg/mL) for one day to induce expression of exogenous *DCLRE1C* cDNA, then transfected by lipofection with *Sp*Cas9-RT (PEnMAX) expressing plasmid and either pegRNA or springRNA expressing plasmid. Overlayed indel distribution patterns (left) and insertion frequencies (right) show results of PEn editing (with pegRNA or springRNA) for *DPM2* target site (11bp intended insertion) in WT or *DCLRE1C* knockout HEK293T cells, with either mRuby or *DCLRE1C* overexpressed. Data analyzed as described in (A) and (B) (n=3 biological replicates ± SD). Intended insertion size on histograms is indicated by a red diamond. *P*-values were determined using t-test (two-tailed, unpaired): * p < 0.05, ** p < 0.01, *** p < 0.001, **** p < 0.0001.

In prime editing, while precise editing is typically the success metric, for certain insertions, stringent precision can be overly restrictive. Longer-than-intended insertions (*e.g.*, protein-epitope tags or integrase attachment sites like *attB/attP*) may be functionally acceptable if the coding sequence is intact or translation is appropriate. This flexibility is crucial for springRNA-mediated PRINS editing, which relies on the inherently less precise NHEJ. In such contexts, prioritizing increased frequency of functionally acceptable insertions (precise and longer-than-intended) can be more beneficial.

Quantitative analysis of indel profiles (categorized into intended insertion size and longer insertions) further supported this (Figure 2b). In the *DCLRE1C* knockout HAP1 cells, the proportional abundance of intended or longer insertions increased by 1.7 to 3.7-fold (springRNA: *DPM2* 13.1% to 48.7%, *PCSK9* 9.4% to 22.6%, *PDCD1* 3.3% to 5.5%, and *TRAC* from 13.3% to 36.2%) compared to wild-type cells, with lesser impact on pegRNA outcomes. For three sites (*PCSK9*, *PDCD1*, *TRAC*), springRNA-mediated insertion frequencies outperformed equivalent pegRNA insertions in the *DCLRE1C* knockout background (pegRNA vs springRNA: *PCSK9* 9.8% vs 22.6%, *PDCD1* 2.1% vs 5.5%, *TRAC* 9.3% vs 36.2%). Even at the *DPM2* target site, where pegRNA yielded quantitatively higher intended or longer insertions (pegRNA vs springRNA *DCLRE1C* knockout: *DPM2* 65.2% vs 48.7%), its performance was predominantly driven by NHEJ-mediated insertion of longer-than-intended products which were improved in the *DCLRE1C* knockout background, highlighting the competitive nature of PEn repair for pegRNAs (involving both homology-driven repair and NHEJ of the 3’-overhang). These observations collectively suggest that springRNA-mediated, NHEJ-driven insertion, especially without Artemis, provides an efficient mechanism for achieving desired insertions, potentially surpassing homology-driven approaches. Thus, Artemis plays a critical role in regulating the processing of PEn-generated 3’-overhangs, and its absence channels repair towards longer insertion outcomes, enhancing PRINS editing efficiency where longer insertions are tolerated.

Beyond Artemis, our initial screen identified *TP53BP1* influencing PEn indel distributions. *TP53BP1* knockout consistently improved pegRNA-mediated editing precision by converting longer-than-intended insertions into more intended ones (Supplementary Figure 4a). For springRNA, *TP53BP1* knockout caused a modest reduction in longer insertions. While *TP53BP1* knockout increased pegRNA intended insertion frequency compared to the wild-type cells (pegRNA: *DPM2* 9.1% to 40.6%, *PCSK9* 3.3% to 9.8%, *PDCD1* 0.3% to 0.7%, *TRAC* 5.4% to 13.0%), it did not increase total intended or longer insertions frequency (Supplementary Figure 4b,c), aligning with findings that 53BP1 inhibition improves pegRNA precision, but not overall editing efficiency [17]. Recently, we also established a strategy (2i+PEn) combining DNA-PKcs and PolΘ inhibitors to improve precision of pegRNA editing. Although these inhibitors could substantially reduce undesired editing by-products of PEn, we could not observe overall improvement in the editing efficiency [18]. This distinction highlights different PEn optimization strategies: *TP53BP1* or DNA-PKcs/PolΘ inhibition would serve more application requiring higher precision, while Artemis absence (with springRNA) offers an advantage for increased overall insertion frequency (including functionally acceptable longer insertions). The choice of DNA repair modulation should align with application requirements for precision versus efficiency.

To further elucidate Artemis’s mechanism, we investigated its interplay with DNA-PKcs. Artemis functions as an NHEJ nuclease recruited and activated by DNA-PKcs [19]. Treating wild-type (WT) and *DCLRE1C* knockout HAP1 cells with a DNA-PKcs inhibitor (DNAPKi) during PEn suppressed all insertions (Figure 2c). For pegRNA, DNAPKi improved precise homology-driven insertion while suppressing other insertions in both WT and *DCLRE1C* knockout cells. Crucially, indel profiles in WT and *DCLRE1C* knockout cells became highly similar with DNAPKi, implying Artemis activity depends on DNA-PKcs.

To establish broader applicability, we assessed *DCLRE1C* knockout effects on springRNA-mediated PRINS editing in other cell types. We observed a similar shift towards longer insertions in HEK293T cells, and to a lesser extent in human induced pluripotent stem cells (hiPSCs), confirming that Artemis’s impact on PRINS editing is not restricted to HAP1 cells (Supplementary Figure 5).

Finally, to directly confirm the phenotype was due to Artemis deficiency, we performed a rescue experiment (Figure 2d). In HEK293T cells (WT and *DCLRE1C* knockout) with a doxycycline-inducible system integrated into the *HBEGF* locus, overexpression of exogenous Artemis in *DCLRE1C* knockout cells reverted the indel profile to resemble wild-type, rescuing the knockout phenotype while mRuby overexpression (control) had no impact. This direct rescue confirms PEn outcome modulation is attributable to Artemis expression levels, confirming its specific role in regulating PEn-generated 3’-overhang processing.

### Length-Dependent Processing of 3’-Overhangs by Artemis Shapes PRINS Editing Outcomes

To define the mechanism for Artemis-dependent processing of 3’-overhangs, we investigated *DCLRE1C* knockout effects across a range of defined insertion lengths. To elucidate specific length requirements for Artemis activity in PEn, we performed PRINS editing at *DPM2* using springRNAs for 1-11 bp insertions (1 bp increments), comparing indel profiles in WT and *DCLRE1C* knockout HEK293T cells (Figure 3a). Our results revealed a clear length-dependent effect: the beneficial shift towards longer insertions in *DCLRE1C* knockout cells only appeared with 6 bp intended insertion (1.2x improvement of intended or longer insertions). This intensified with increasing length, becoming most pronounced and significant for 8 bp intended insertion and longer (over 2X improvement of intended or longer insertions). For intended insertions ≥ 8 bp, the *DCLRE1C* knockout background consistently yielded a higher proportion of intended or longer insertions. This indicates that Artemis primarily engages with and processes 3’-overhangs of sufficient minimum length, and that its impact on shaping indel outcomes increases with reverse-transcribed overhang length.

**Figure 3.**
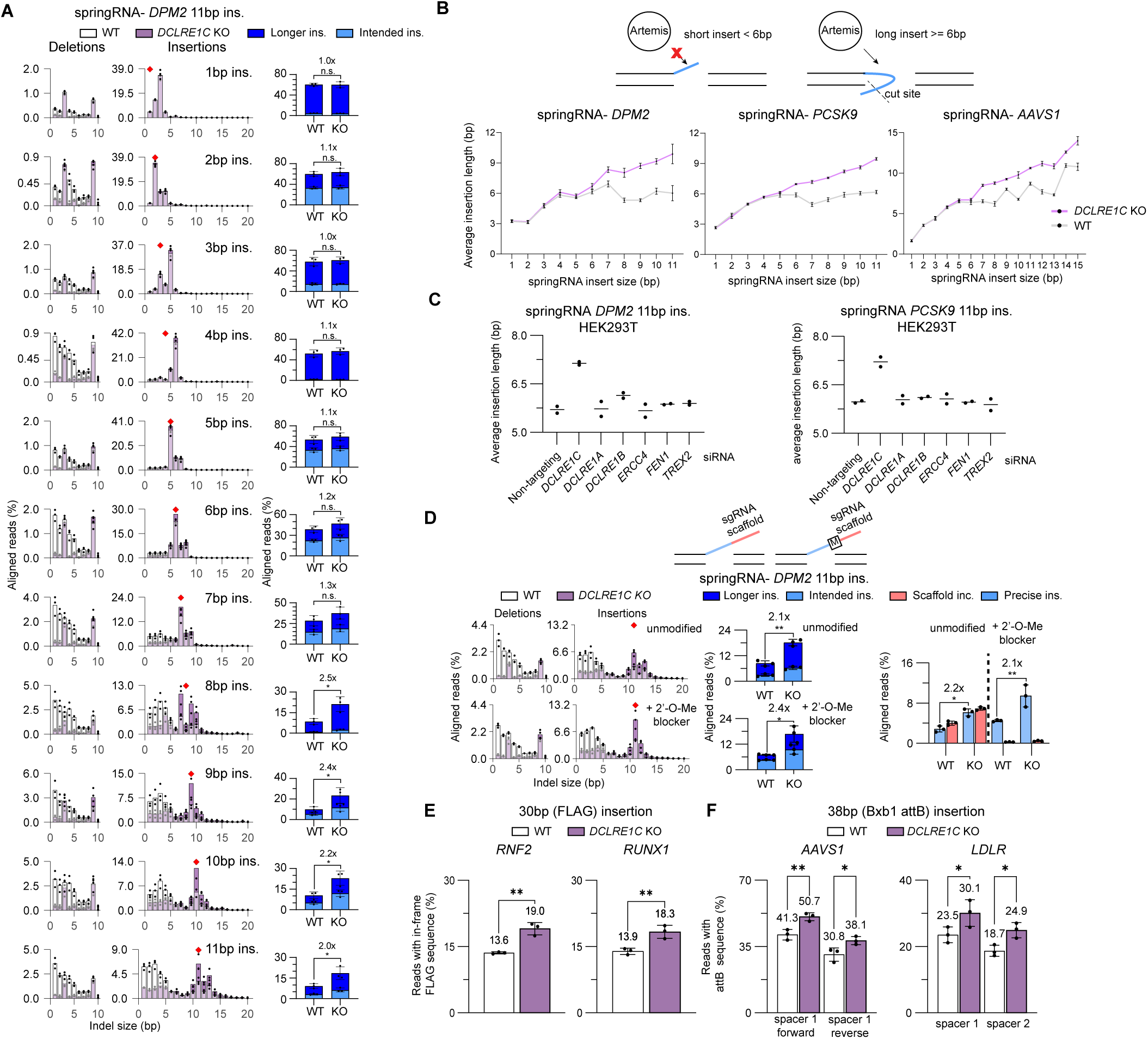
Length-Dependent Processing of 3’-Overhangs by Artemis Shapes PRINS Editing Outcomes. A) Indel distribution and insertion frequencies for PEn editing with springRNA (varying sizes) in WT and *DCLRE1C* knockout HEK293T cells. Cells were transfected with PEnMAX plasmid and synthetic springRNA (1-11bp insertions) at the *DPM2* target site. Overlayed indel histograms (CRISPResso2, custom R-script) show average indel size frequencies (three biological replicates) for WT (white) and *DCLRE1C* knockout (purple) at *DPM2* for indicated insertion sizes. Bar graph shows average frequency (n=3 biological replicates ± SD) of intended or longer insertions calculated using CRISPResso2 and a custom R-script. Intended insertion size is indicated by a red diamond. B) Average insertion length of springRNA (increasing insert size) in WT and *DCLRE1C* knockout HEK293T cells. Schematic (top) illustrates effect of Artemis on different insert lengths, with minimal effect on shorter overhangs (<6 bp) and more pronounced effect on longer overhangs (≥ 6bp). Cells were transfected with PEn plasmid and synthetic springRNAs for *DPM2* (1-11bp), *PCSK9* (1-11bp), and AAVS1 (*PPP1R12C*) (1-15bp) sites. Average insertion length calculated as weighted frequency of each length divided by sum of all insertion frequencies (n=3 biological replicates ± SD) (CRISPResso2, custom R-script). C) siRNA-mediated knockdown of Artemis (*DCLRE1C*), but not other nucleases, uniquely increases average insertion length of PRINS. HEK293T cells were subjected to siRNA-mediated knockdown of either Artemis (*DCLRE1C*), or other known nucleases, alongside non-targeting siRNA control. Following knockdown, cells were transfected with PEnMAX plasmid and springRNA plasmids designed for 11 bp insertions at two distinct genomic loci, *DPM2* and *PCSK9*. Average insertion length calculated as weighted frequency of each length divided by sum of all insertion frequencies (n=2 biological replicates) (CRISPResso2, custom R-script). D) Indel distribution patterns and editing outcomes for PRINS editing using either a standard springRNA or a 2’-O-methylation scaffold modified (2’-O-Me blocker) springRNA in WT and *DCLRE1C* knockout HEK293T cells. Cells were transfected with PEnMAX plasmid and synthetic springRNA (11 bp insertion) at the *DPM2* target site either with or without the scaffold modification. Overlayed indel histograms (CRISPResso2, custom R-script) show average indel size frequencies (n=3 biological replicates) comparing WT (white) and *DCLRE1C* knockout (purple) backgrounds for standard springRNA (left-above) and scaffold modified (left-below) alongside insertion frequencies bar graphs. Intended insertion size is indicated by a red diamond. Bar graphs (right) show average frequencies (n=3 biological replicates ± SD) of’Precise ins’ (unmodified prime edits) and’Scaffold inc.’ (all scaffold incorporations) for unmodified springRNA (right-above) and 2’-O-methylated scaffold springRNA (right-below), quantified by amplicon-seq and CRISPResso2. E) FLAG tag (30 bp) insertion rate in WT and *DCLRE1C* knockout HEK293T cells. Cells were transfected with PEnMAX plasmid and springRNA plasmids for FLAG insertion at two target sites (*RNF2*, *RUNX1*). Percentage of in-frame FLAG tag insertions (intended or longer) analyzed by amplicon-seq (CRISPResso2, custom R-script), counting reads with intended insertion or longer with no other unintended indels 10 bp downstream of the cut site. Bar graph shows average percentage (n=3 biological replicates ± SD) of in-frame FLAG insertions. F) Bxb1 *attB* (38bp) insertion rate in WT and *DCLRE1C* knockout HEK293T cells. Cells were transfected with PEnMAX plasmid and springRNA plasmids for *attB* insertion at two target sites (AAVS1 (*PPP1R12C*), *LDLR*). Percentage of *attB* site insertions analyzed by amplicon-seq (CRISPResso2, custom R-script), counting reads with intended or longer insertion. Bar graph shows average percentage (n=3 biological replicates ± SD) of *attB* insertions. *P*-values were determined using t-test (two-tailed, unpaired): * p < 0.05, ** p < 0.01, *** p < 0.001, **** p < 0.0001. Ins. = insertions; WT = wild-type; KO = *DCLRE1C* knockout.

To further quantify this, we calculated the average insertion length for springRNA-mediated editing at three different target sites (*DPM2*, *PCSK9*, and AAVS1 (*PPP1R12C*)) across programmed insertion ranges (*DPM2*: 1-11 bp; *PCSK9*: 1-11 bp; AAVS1 (*PPP1R12C*): 1-15 bp) (Figure 3b). The average insertion length was calculated by dividing the weighted frequency of each insertion length by the sum of all insertion frequencies. Consistent with indel profiles, the average insertion length in *DCLRE1C* knockout cells diverged from wild-type only when intended insertions reached 6-7 bp. For longer intended insertions, average length in the knockout was for the most part higher, reinforcing that Artemis’s processing activity is 3’-overhang length-dependent.

These cellular data are consistent with *in vitro* biochemical studies: Artemis, activated by DNA-PKcs, functions as both an endonuclease and exonuclease processing 3’-DNA overhangs, proposing it trims longer 3’-overhangs by folding it into a hairpin-like structure before cleaving, leaving a residual 4 bp 3’-overhangs [20, 21]. Our data strongly supports this model, suggesting that in Artemis’s absence, normally resected longer 3’ overhangs are instead retained and integrated via NHEJ, causing the observed shift towards longer and more frequent insertions in PRINS editing. We presumed the effect by Artemis knockout would likely be unique amongst other nucleases within its gene family because it has both endonuclease activity and exonuclease activity to process 3’-overhangs [21]. We therefore also tested the effect of siRNA knockdown on Artemis (*DCLRE1C*) and other nucleases, including *DCLRE1A* and *DCLRE1B*, on the average insertion length of 11 bp inserts into *DPM2* and *PCSK9* (Figure 3c). Only targeted knockdown of Artemis gene (*DCLRE1C*), and not any other nuclease, led to a substantial increase in the average insertion length supporting the case that this is a unique role of Artemis.

Given PEn’s propensity (with springRNAs) to incorporate the longer-than-intended insertion that includes the guide RNA scaffold, we explored mitigation strategies. Our previous work demonstrated that internal chemical modification of the pegRNA/springRNA with different chemical modifiers, such as 2’-O-methylation, blocks reverse transcriptase read-through and prevents scaffold incorporation [22]. We investigated the combined effect of this modification and Artemis knockout on PRINS editing outcomes at *DPM2* (Figure 3d). Indel profiles revealed the 2’-O-methylation blocker reduced longer-than-intended insertions, but not completely. Importantly, even with the 2’-O-methylation blocker, the beneficial Artemis knockout effect on indel distribution (shift towards longer insertions) persisted. A closer look at the categorization of insertions (Figure 3d, right panel) confirmed the 2’-O-methylation blocker largely eliminated scaffold-incorporated reads. Remaining longer-than-intended insertions were primarily single nucleotide, non-templated additions (*e.g*., T or C), distinct from scaffold-derived insertions seen with unmodified springRNAs (Supplementary Figure 6). With normal springRNA, Artemis knockout improved both precise and scaffold-incorporated reads. Combined with chemically modified springRNA, Artemis knockout specifically led to more precise insertions, while retaining the overall shift towards longer desired insertions. This demonstrates that combining Artemis knockout with the chemical modifications increases the frequency of precise insertions, offering a useful strategy for applications where strict precision is desired within high efficiency.

To illustrate practical utility, we investigated if enhanced PRINS insertion in an Artemis-deficient background could be leveraged for gene editing. We tested FLAG protein epitope tag (30 bp) insertion via PRINS editing at two different endogenous loci previously tested [23]: *RNF2* and *RUNX1* (Figure 3e). In both cases, Artemis (*DCLRE1C*) knockout improved overall FLAG tag insertion frequency of either intended or longer insertions from 13.6% to 19.0% (*RNF2)* and 13.9% to 18.3% (*RUNX1)*.

We also evaluated Bxb1 serine integrase *attB* attachment site (38 bp) insertion, used for large cargo integration with serine integrases [24], at two distinct genomic target sites, utilizing either two different directions of insertion or two different spacers (Figure 3f). Across all four target-spacer combinations, Artemis (*DCLRE1C*) knockout consistently enhanced overall *attB* insertion frequency by 5% to 10% of total edited reads (attB insertion: AAVS1 (*PPP1R12C*) spacer 1 (insert forward direction) 41.3% to 50.7%, AAVS1 (*PPP1R12C*) spacer 1 (insert reverse direction) 30.8% to 38.1%, *LDLR* intron 1 spacer 1 23.5% to 30.1%, *LDLR* intron 1 spacer 2 18.7% to 24.9%). These results demonstrate that the absence of Artemis provides a highly effective strategy to enhance PRINS editing efficiency for larger functional sequences, offering substantial advancement for diverse genomic engineering applications.

### Modulations of Artemis Expression Recapitulate Beneficial PRINS Editing Outcomes

Artemis deficiency enhances PRINS editing efficiency for longer insertions, and this effect can be combined with gRNA modifications for tailored precision. Translating these insights often requires transient, controllable gene expression modulation. Therefore, we investigated epigenetic silencing (CRISPRoff), siRNA, and antisense oligonucleotides (ASOs) to reduce Artemis expression and recapitulate the beneficial PEn outcomes of *DCLRE1C* knockout.

We evaluated knockdown efficiency and duration in HEK293T cells over seven days (Figure 4a). For epigenetic silencing, we employed the CRISPRoff system with deactivated *Sp*Cas9 (D10A,H840A) fused to a KRAB domain and methyltransferase domains (*DNMT3A/3L*) [25], guided by a gRNA targeting the *DCLRE1C* promoter [26]. This method achieved >90% Artemis knockdown compared to the non-targeting gRNA, sustained from day 1 to day 7, including one cell passage. In contrast, siRNA-mediated knockdown (four-siRNA pool targeting *DCLRE1C*) yielded less robust silencing (<50% KD on day 1), with Artemis expression recovering after one passage. We initially screened different ASO designs to target *DCLRE1C* transcript and used the best candidate for comparative analysis of knockdown efficiency (Supplementary Figure 7). ASO-mediated knockdown achieved comparable KD efficiency to epigenetic silencing (>90% on day 1); however, like siRNA, expression levels rapidly recovered after one passage. These results highlight distinct knockdown kinetics and durations and aid selection of appropriate transient modulation strategies.

**Figure 4.**
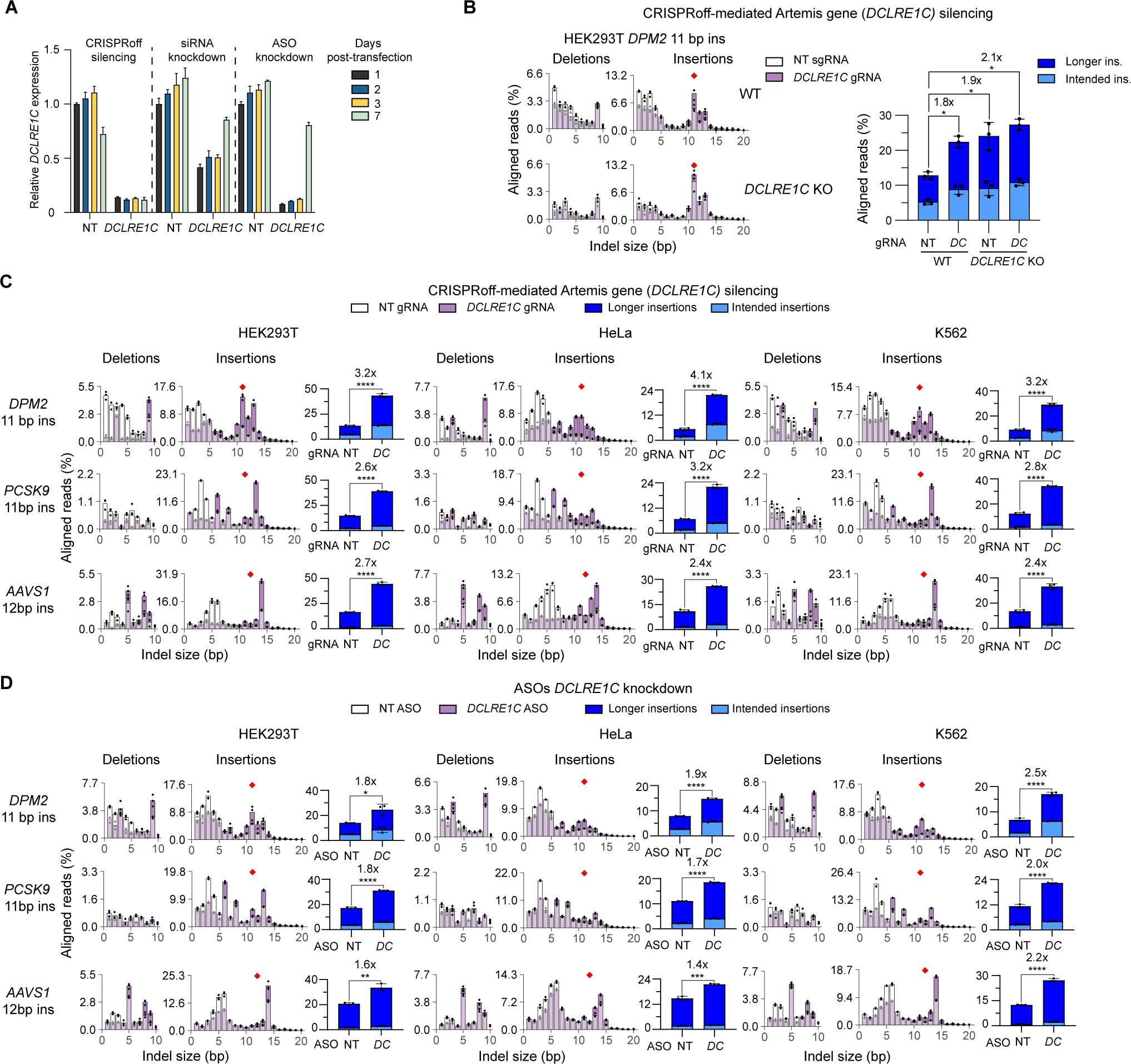
Transient Modulations of Artemis Expression Recapitulate Beneficial PRINS Editing Outcomes A) Comparison of Artemis gene *(DCLRE1C)* knockdown/silencing efficiency in HEK293T cells by CRISPRoff, siRNA, and antisense oliginucleotides (ASOs). Cells were transfected (lipofection) with either *DCLRE1C*-targeting or non-targeting CRISPRoff components (CRISPRoff mRNA + gRNA), siRNA, or ASO. *DCLRE1C* expression was measured by qPCR at indicated time points (Day 1, 2, 3, 7) using the appropriate non-targeting condition as the control group and *TBP* as the reference gene. Bar graph shows average fold expression change of *DCLRE1C* relative to Day 1 non-targeting conditions for each modality (n=3 biological replicates ± SD). B) Effect of CRISPRoff-mediated Artemis (*DCLRE1C*) gene silencing on PRINS-mediated insertion in WT and *DCLRE1C* knockout HEK293T cells. Both cell lines were transfected (lipofection) with CRISPRoff components (CRISPRoff mRNA + Artemis-targeting or non-targeting gRNA), grown and maintained for at least 5 days, then transfected with PEnMAX plasmid and springRNA plasmid for *DPM2* 11 bp insertion. Overlayed indel histograms (CRISPResso2, custom R-script) show average indel frequencies (n=3) for non-targeting (white) and *DCLRE1C*-targeting gRNA (purple) conditions within each cell background. Intended insertion size on histograms is indicated by a red diamond. Bar graph shows average frequency (n=3 biological replicates ± SD) of intended or longer insertions calculated using CRISPResso2 and a custom R-script. NT = non-targeting gRNA; *DC* = *DCLRE1C* promoter targeting gRNA. Ins. = insertions. C) CRISPRoff-mediated Artemis gene (*DCLRE1C*) silencing for PRINS insertion in different cell types. HEK293T, HeLa, and K562 cells were electroporated with CRISPRoff components (*DCLRE1C*-targeting or non-targeting gRNA), grown and maintained for at least 5 days, then electroporated with PEnMAX mRNA, La (1-194) mRNA, and synthetic springRNAs for *DPM2*, *PCSK9*, and AAVS1 (*PPP1R12C*) insertions. Overlayed indel histograms (CRISPResso2, custom R-script) show average indel frequencies (n=3) for non-targeting (white) and *DCLRE1C*-targeting gRNA (purple) conditions within each cell background. Intended insertion size on histograms is indicated by a red diamond. Bar graph shows average frequency (n=3 biological replicates ± SD) of intended or longer insertions across all cell types and targets calculated using CRISPResso2 and a custom R-script. NT = non-targeting gRNA; *DC* = *DCLRE1C* promoter targeting gRNA. D) Effect of ASO-mediated Artemis gene (*DCLRE1C*) knockdown on PRINS-mediated insertion in HEK293T, HeLa, and K562 cells. Cell lines were electroporated with *DCLRE1C*-targeting or non-targeting ASOs for 3 days, then electroporated with PEnMAX mRNA, La (1-194) mRNA, and springRNAs for *DPM2*, *PCSK9*, and AAVS1 (*PPP1R12C*) insertions. Overlayed indel histograms (CRISPResso2, custom R-script) show average indel frequencies (n=3) comparing non-targeting (white) and *DCLRE1C*-targeting ASOs (purple) conditions. Intended insertion size on histograms is indicated by a red diamond. Bar graph shows average frequency (n=3 biological replicates ± SD) of intended or longer insertions calculated using CRISPResso2 and a custom R-script. NT = non-targeting ASO; *DC* = *DCLRE1C* promoter targeting ASO. *P*-values from Figure 4b were determined using ANOVA one-way test (two-tailed, unpaired) with multiple comparisons corrected using Tukey’s test. Displayed pairwise comparisons are relative to NT gRNA WT background. *P*-values from Figure 4 c, d were determined using t-test (two-tailed, unpaired): * p < 0.05, ** p < 0.01, *** p < 0.001, **** p < 0.0001.

Next, we investigated whether epigenetic silencing of Artemis could recapitulate beneficial PRINS editing outcomes in HEK293T cells. We transfected WT and *DCLRE1C* knockout cells with the CRISPRoff editor (mRNA) along with either a non-targeting guide RNA or *DCLRE1C*-targeting guide RNA. Following one passage, these cells were then transfected with the PEn editor and a springRNA targeting a *DPM2* site (Figure 4b). In WT cells, epigenetic silencing of *DCLRE1C* resulted in a characteristic indel shift similar to *DCLRE1C* knockout cells treated with either non-targeting or *DCLRE1C*-targeting silencing gRNA, improving intended or longer insertions by 1.8-fold. These results confirm epigenetic silencing effectively mimics the Artemis knockout phenotype.

To assess the broader applicability of epigenetic silencing of Artemis and PRINS, we extended our analysis to different cell lines using all-RNA components. We performed Artemis gene (*DCLRE1C*) silencing using the CRISPRoff components, and after one passage, delivered PEn mRNA and synthetic springRNAs targeting *DPM2* (11 bp), *PCSK9* (11 bp), and AAVS1 (*PPP1R12C*) (12 bp) (Figure 4c). Also to improve overall editing efficiency, we added to each conditions mRNA overexpressing truncated La protein and springRNA with La-accessible polyU tails, which stabilizes and protects the springRNA, as was describe with prime editing PE7 [27]. Additionally, for consistency in delivery and for overall improvement in silencing/prime editing efficiency, all components were delivered to the cells by electroporation instead of lipid-based transfection methods. Silencing efficiency, confirmed by qPCR, was high across all cell lines, ranging from over 99% in HEK293T and HeLa cells, to 95% in K562, and approximately 90% in HepG2 cells (Supplementary Figure 8a).

Intriguingly, analysis of relative Ct values suggested that HEK293T and K562 cells exhibited the highest basal expression levels of Artemis, followed by HeLa, with HepG2 showing the lowest endogenous expression.

Analysis of PRINS editing outcomes revealed that Artemis silencing generally shifted indel profiles and increased intended or longer insertions (Figure 4c, Supplementary Figure 8b). The effect was comparable between HEK293T, HeLa, and K562 cells, where improvement in intended or longer insertions reached 2-4-fold across tested sites, with indel distribution plots showing visibly higher peaks corresponding to longer insertions. However, in HepG2 cells, the beneficial effect of Artemis silencing was less pronounced (Supplementary Figure 8b). This differential response, where HepG2 (with the lowest basal Artemis expression) showed a muted improvement compared to the other cell lines (which had higher basal Artemis expression), suggests that the beneficial effect of Artemis silencing on PRINS editing efficiency is more significant in cell types with higher endogenous Artemis expression. This implies a context-dependent impact, potentially reflecting a specialized role of Artemis in these lineages or a threshold effect related to its basal expression level.

Beyond epigenetic silencing, we also investigated the use of ASOs as another transient method to modulate Artemis expression across HEK293T, HeLa, and K562. Delivery methods (electroporation) were kept the same as with epigenetic silencing to be consistent between cell types. Knockdown efficiency and duration were evaluated by qPCR over a time course (Supplementary Figure 9). Close to 90% knockdown efficiency was achieved in all cell lines after 2 days, but expression levels started recovering after only three days and were completely restored after 9 days. In all three cell lines, ASO-mediated *DCLRE1C* knockdown led to a general improvement in PRINS editing with all-RNA components at the three target sites (*DPM2*, *PCSK9*, and AAVS1(*PPP1R12C*)) including the characteristic indel shift and significant increase in intended or longer reads, but not to the same extent as the epigenetic silencing (Figure 4d). These findings validate ASOs as a transient strategy for Artemis modulation to improve PRINS insertions but will not completely replicate the effect of the knockout or epigenetic silencing.

The successful demonstration of both epigenetic silencing and ASO-mediated strategies for reducing Artemis expression, and their ability to recapitulate the beneficial PRINS editing outcomes of Artemis knockout cells, enables tools for improved control over insertion efficiency. These modulation methods allow for fine-tuning of PRINS editing outcomes non-permanently, opening diverse avenues for advanced gene editing applications. This data also suggests that PRINS based assays can be leveraged to discover Artemis inhibitors that may have therapeutic value, particularly in oncology settings.

### Artemis Modulation to Improve PRINS-Mediated Endogenous Protein Tagging

Leveraging our comprehensive understanding of Artemis’s role in PRINS editing and the development of effective transient knockdown strategies, we next explored the utility of modulating Artemis for functional gene editing applications.

We set out to illustrate the use of PRINS editing for endogenous protein tagging (Figure 5a). This involves inserting a sequence encoding an epitope tag (*e.g.*, HIBIT, FLAG) at the N- or C-terminus of an endogenous gene with an appropriate short linker peptide, allowing for functional protein labelling under its native promoter. To evaluate the baseline efficiency of PRINS-mediated tagging, we inserted a HIBIT tag [28] at the C-terminus of *GAPDH* in HEK293T cells using a springRNA (Figure 5b). HIBIT activity, indicative of successful in-frame tagging, was detected only when both springRNA and *Sp*Cas9-MMLV RT components were expressed. Notably, the tethered PEnMAX format yielded higher HIBIT activity compared to untethered *Sp*Cas9 and MMLV RT, confirming PEnMAX’s superior performance for this application. We further validated the applicability of PRINS tagging in a more challenging conditions using all-RNA components, demonstrating HIBIT activity for *GAPDH* C-terminal tagging in primary human hepatocytes exclusively under PEnMAX mRNA and synthetic springRNA conditions (Figure 5b).

**Figure 5.**
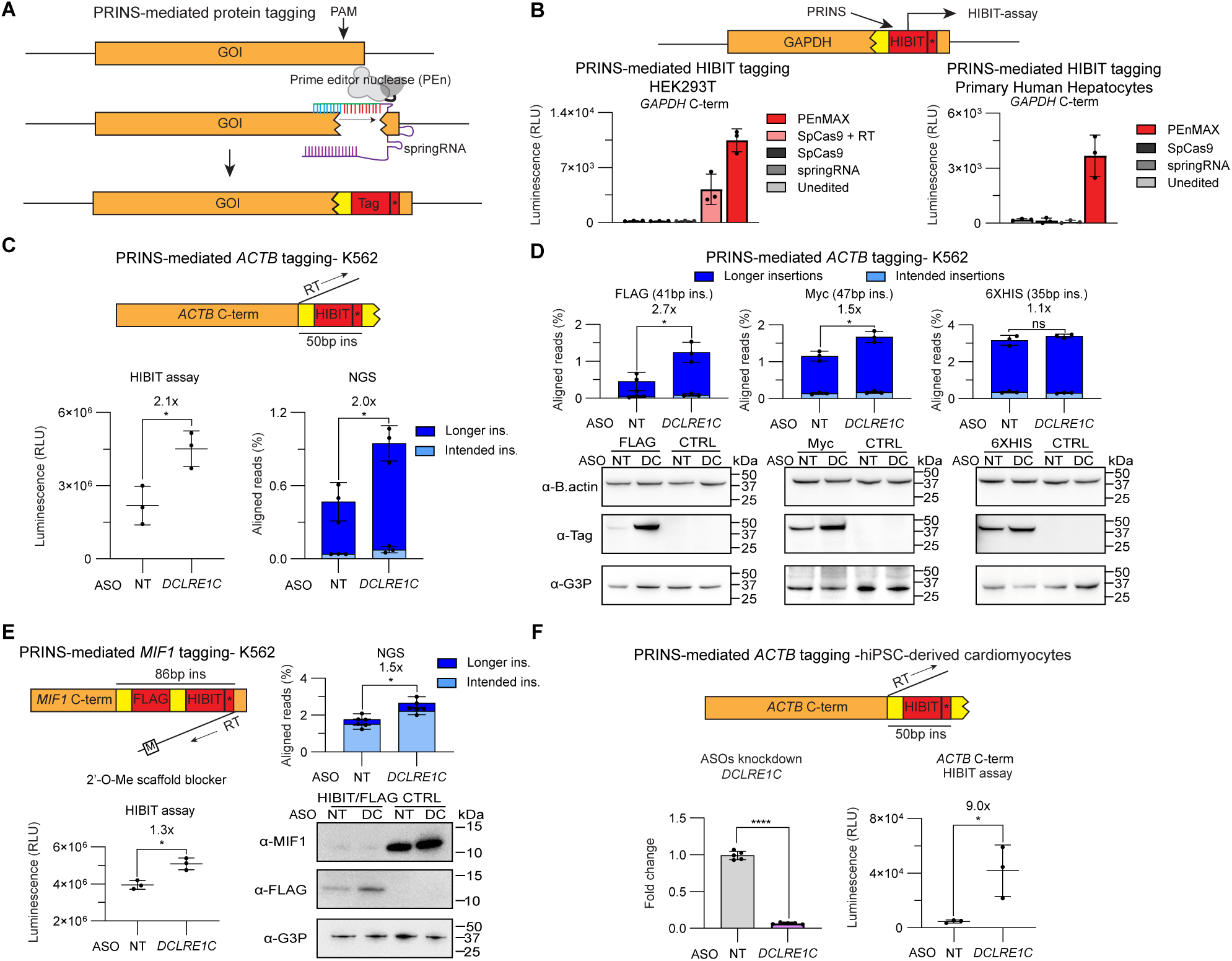
Artemis Knockdown Improves PRINS-mediated Endogenous Protein Tagging A) Schematic representation of PRINS-mediated endogenous protein tagging. Protospacer adjacent motif (PAM) sites for PRINS editing are chosen near the terminal end of a gene of interest (GOI) to insert a desired protein tag and associated linker in-frame, including additional start/stop codons. B) PRINS-mediated HIBIT tagging of endogenous *GAPDH* on the C-terminal end in HEK293T and primary human hepatocytes (PHHs). HEK293T cells were transfected with synthetic springRNA for GAPDH C-terminal HIBIT tagging, alongside various PEn component combinations (*Sp*Cas9, untethered M-MLV RT, PEnMAX, or controls) as plasmids. PHHs were similarly transfected with springRNA and PEn components (*Sp*Cas9, PEnMAX, or controls) as mRNA. HIBIT activity, indicative of successful in-frame tagging, was assessed via Promega HIBIT lytic assay (average luminescence signal, n=3 biological replicates ± SD) 2-3 days post-transfection. C) Effect of ASO-mediated Artemis gene (*DCLRE1C*) knockdown on PRINS-HIBIT tagging on *ACTB* C-term in K562 cells. Cells were electroporated with *DCLRE1C*-targeting or non-targeting ASOs, then electroporated with PEnMAX mRNA, La (1-194) mRNA, and springRNAs for *ACTB* C-term HIBIT tagging (50 bp insertion). Dot plots (left) display average HIBIT luminescence (n=3 biological replicates ± SD) for HIBIT-tagged samples. Bar graphs (middle) show average frequency (CRISPResso2, n=3 biological replicates ± SD) of intended or longer insertions. D) Effect of ASO-mediated Artemis gene (*DCLRE1C*) knockdown on different protein tags. Cells were treated as in c) but with springRNA for *ACTB* C-term FLAG, MYC, and poly-histidine tagging (47, 41, 35 bp insertions, respectively). Bar graphs (top) show average frequency (CRISPResso2, n=3 biological replicates ± SD) of intended or longer insertions at each target site. Western blot images (right) illustrate protein tagging efficiency for selected tags on the beta actin protein under *DCLRE1C* knockdown, using specific tag antibodies, beta actin antibodies, and GAPDH (G-3-P) as a loading control. E) Effect of ASO-mediated Artemis gene (*DCLRE1C*) knockdown on longer PRINS-mediated insertion. Cells were treated as in c) but with 2’-O-methylated springRNA for *MIF1* C-term dual FLAG and HIBIT tagging (86bp insertions). Dot plots (left) display average HIBIT luminescence (n=3 ± SD), bar graphs show average frequency (n=3 biological replicates ± SD) of intended or longer insertions and Western blot images illustrate MIF1 FLAG protein tagging efficiency under *DCLRE1C* knockdown. F) Effect of ASO-mediated Artemis gene (*DCLRE1C*) knockdown on PRINS-mediated endogenous protein tagging in hiPSC-derived cardiomyocytes cells. Cells were transfected (lipofection) with *DCLRE1C*-targeting or non-targeting ASOs, then transfected (lipofection) with PEnMAX mRNA and springRNAs for *ACTB* C-term HIBIT tagging. Bar graph (left) shows average ASO-mediated *DCLRE1C* knockdown efficiency (n=5 biological replicates ± SD). Dot plots (right) display average HIBIT luminescence (n=3 biological replicates ± SD) for HIBIT-tagged samples following ASO-mediated *DCLRE1C* knockdown in separately transfected cells. NT = non-targeting ASO, DC = *DCLRE1C*-targeting ASO. *P*-values from Figure 5 were determined using t-test (two-tailed, unpaired): * p < 0.05, ** p < 0.01, *** p < 0.001, **** p < 0.0001.

Having established robust PRINS tagging, we next evaluated endogenous protein tagging under transient modulation of Artemis using antisense oligonucleotides (ASOs). While epigenetic silencing (e.g., CRISPRoff) can provide more stable—and in some contexts more pronounced—repression, our aim was to illustrate an approach suited to tagging workflows that require only a short editing window. ASO-mediated knockdown temporarily decreases *DCLRE1C* expression to boost PRINS insertion efficiency, then permits rapid recovery of Artemis levels after the editing phase, thereby avoiding sustained alteration of DNA repair pathways. Thus, although epigenetic silencing may deliver more durable effects, ASOs are better aligned to protein tagging applications where transient modulation is desirable.

We designed a springRNA to tag the C-terminal end of the highly expressed beta-actin gene (*ACTB*) with HIBIT and a short linker, followed by a stop codon to terminate translation (50 bp total insertion). We tested this springRNA in ASO-treated K562 cells using all-RNA components, consisting of PEnMAX mRNA, truncated La mRNA, and synthetic springRNA with La-accessible polyU tail (Figure 5c). ASO-mediated *DCLRE1C* knockdown significantly improved HIBIT activity for *ACTB* tagging, increasing levels of HIBIT activity in the cell extract by 2.1-fold. The relative activity levels correlated well with NGS data showing increased frequency of intended or longer insertions in cells treated with *DCLRE1C*-targeting ASO (2.0-fold). To demonstrate versatility, we further evaluated PRINS-mediated protein tagging efficiency but with FLAG, MYC, and poly-histidine tags (Figure 5d). Corresponding NGS results confirmed, for the most part, increased frequency of intended or longer insertions for each tag under *DCLRE1C* knockdown, although to different degrees (Figure 5d, top). FLAG tagging and Myc tagging efficiency was improved 2.7-fold and 1.5-fold, respectively, but poly-histidine tagging was not significantly improved. The differing outcomes between the different tags in response to ASO-mediated *DCLRE1C* knockdown suggest that the sequence context (e.g. G/C content) and the precise nature of the 3’-generated overhang can affect insertion efficiency, potentially modulating or even obscuring the beneficial effects of Artemis knockdown. Western blot analysis further correlated with the NGS data, with some conditions showing higher levels of tagged proteins when *DCLRE1C* knockdown was applied (Figure 5d, bottom). Consistent with the NGS data, Western blot for the poly-histidine tag at *ACTB* showed no change in tagging efficiency between non-targeting and *DCLRE1C*-targeting ASO conditions. But for FLAG and Myc tags, the tagged beta-actin protein was visibly higher in the presence of *DCLRE1C* knockdown compared to the control conditions, while total beta actin levels remained constant. These show that Artemis knockdown can in certain situations (e.g. FLAG tagging *ACTB*) increase the level of PRINS-mediated protein tagging to measurable degrees.

To further probe the capabilities of PRINS-mediated protein tagging and assess its size limit, we also attempted to insert a longer tag containing both the FLAG tag and the HIBIT tag, separated by short linkers (86 bp total insertion) on the C-terminus of the macrophage migration inhibitor factor gene (*MIF1*). Orientation of the optimal spacer for this target was such that longer-than-intended insertions would create out-of-frame insertions. Therefore, we applied 2’-O-methylation modification on the scaffold of the springRNA to mitigate these longer-than-intended insertion events. For this dual tag insertion, ASO-mediated *DCLRE1C* knockdown improved HIBIT activity compared to the non-targeting control by 1.3-fold, which correlated well with NGS data (1.5-fold increase in intended or longer insertions) (Figure 5e, bottom left and top right). Corresponding Western blot data show that overall MIF protein levels were dramatically reduced after PRINS editing compared to untreated control conditions likely caused by the generation of coding-sequence disrupting indels. However, FLAG-tagged MIF1 was detected at the expected protein size (12.5 kDa), and FLAG-tagged levels were visibly increased in cells treated with *DCLRE1C*-targeting ASO compared to non-targeting ASO control (Figure 5e, bottom right).

Crucially, to demonstrate the utility of Artemis modulation in challenging and therapeutically relevant non-dividing cells, we applied ASO-mediated *DCLRE1C* knockdown in hiPSC-derived cardiomyocytes (Figure 5f, left). We achieved a high knockdown efficiency of 95% as confirmed by qPCR. Leveraging this repression, we performed PRINS-mediated HIBIT tagging of the endogenous *ACTB* gene.

Notably, *DCLRE1C* knockdown led to a 9-fold improvement in HIBIT activity compared to non-targeting controls (Figure 5f, right). This substantial enhancement in a terminally differentiated, non-dividing cell type underscores the practical utility and broad applicability of Artemis-modulated PRINS editing, as such cells are notoriously difficult to engineer with traditional homology-dependent methods due to their reliance on NHEJ and inherently low HDR activity. This finding strongly supports the application of Artemis-modulated PRINS editing for gene tagging or modification in various cell types, including difficult-to-transfect and non-proliferating tissues, where current editing strategies face significant hurdles.

## Discussion

Our study identifies Artemis (*DCLRE1C*) as a key regulator of nuclease-based prime editing (PEn) outcomes, specifically demonstrating its capacity to shape indel profiles generated during springRNA-mediated PRINS editing. This discovery not only uncovers a novel mechanism to enhance insertion efficiency in prime editing but also provides in cell insights into the function of Artemis in processing 3’- DNA overhangs of sufficient length.

This nuanced understanding of Artemis’s length-dependent activity also clarifies why its role may have been overlooked in other CRISPR-based genetic screens. Unlike PEn, many CRISPR screens focus on repair outcomes from simpler DSBs or nick-based systems, or on smaller indels, where the generation of long 3’-overhangs—the specific substrate for Artemis’s unique processing activity—is not a predominant feature. Our findings thus highlight the importance of considering the specific nature of DNA repair intermediates when evaluating the contribution of different repair factors.

Furthermore, the unique ability of PRINS editing to generate 3’-overhangs of defined, variable lengths makes it a useful tool for studying diverse DNA repair mechanisms in cells.

Modulating Artemis by epigenetic silencing or ASO recapitulates the knockout phenotype and provides a practical, tunable lever to increase insertion frequency. Crucially, this is exemplified by the robust ∼9-fold enhancement of HIBIT tagging observed in iPSC-derived non-dividing cardiomyocytes upon Artemis knockdown (Figure 5f). These terminally differentiated cells are known for their high reliance on NHEJ and low HDR activity, making them particularly challenging for traditional gene editing. The ability to achieve significant functional editing improvements in such a non-dividing, primary-like cell type demonstrates the potential of Artemis-modulated PRINS editing for applications in post-mitotic tissues, opening new avenues for therapeutic interventions where other methods fall short. However, limitations remain: Outcomes vary by target site and sequence context; relative gains by Artemis knockdown diminish for very long inserts; and synthetic springRNAs can be less efficient than plasmid expression. Continued optimization of editor processivity, RNA stability, and target-site selection should extend performance across additional loci and cargo sizes.

Beyond the functional tagging of endogenous genes, this enhanced PRINS editing platform holds potential for other genomic interventions. For instance, it could be leveraged for the efficient insertion of transcription factor binding sites (TFBS) into promoter or enhancer regions to modulate gene expression [29]. Importantly, PRINS can be adapted for targeted mutagenesis screens to rapidly evaluate protein variants with enhanced function or to introduce synthetic or natural DNA-binding sites that fine-tune transcriptional output. This mirrors the natural diversification achieved by V(D)J recombination, which generates functional variation through programmed DNA rearrangements and produces far more variants than classical point mutagenesis. In addition, an Artemis pulldown–based profiling workflow—similar to DISCOVER-Seq [30]—could be used to capture and sequence repair-associated DNA ends, enabling detection of off-target events from prime editors. The inherent flexibility of PRINS editing, coupled with the tuneable modulation of Artemis, provides a toolset for researchers to insert arbitrary DNA sequences into specific genomic loci.

## Methods

### Ethical Statement

AstraZeneca has a governance framework and processes in place to ensure that commercial sources have appropriate patient consent and ethical approval in place for collection of the samples for research purposes including use by for-profit companies.

### Protein, DNA, mRNA, guideRNAs

Recombinant PEn protein was expressed and purified, as described in [22]. Plasmid expressed PEnMAX, the nuclease version of the PEMAX construct [11], was generated by gene synthesis and cloned into expression vector containing CMV promoter (Genscript). Capped PEnMAX, La (1-194), and *Sp*Cas9-based CRISPRoff editor mRNA was synthesized through T7-directed *in vitro* transcription using a linearized DNA template. Translated sequences of PEnMAX, La (1-194), and CRISPRoff editor are listed in Supplementary Data 1.

Plasmid expressed pegRNAs or springRNAs were generated by gene synthesis and cloned into the pMA vector together with an upstream U6 promoter and an additional‘G’ on the 5’ end of the spacer, if required (Invitrogen). Synthetic pegRNAs, springRNAs, or gRNAs (< 150nt) were chemically synthesized with appropriate end-protecting chemical modifications by either IDT or Genscript. Other springRNA (> 150nt) were special ordered through Genscript. All pegRNA, springRNA, and gRNA sequences, including chemical modifications, used in this study are listed in Supplementary Data 2.

### Small Molecule Compounds, siRNA, ASOs

DNA-PKcs inhibitor AZD7648 was purchased from MedChemExpress (HY-111783) and dissolved in dimethyl sulfoxide (DMSO) at a stock concentration of 10 mM. Non-targeting (D-001810-10-05), *DCLRE1C*-targeting (L-004269-00-0005), *DCLRE1A*-targeting (L-010790-00-0005), *DCLRE1B*-targeting (L-015780-00-0005), *ERCC4*-targeting (L-019946-00-0005), *FEN1*-targeting (L-010344-00-0005), and *TREX2*-targeting (L-032280-00-0005) siRNA pools were purchased from a commercial vendor (Dharmacon), and were all resuspended in RNAse free water at a stock concentration of 100 µM. ASOs in Supplementary Figure 7 were designed by the Data Sciences and Quantitative Biology team and synthesized by the Oligonucleotide Chemistry and Targeted Delivery team, both at AstraZeneca, Sweden. Purified ASOs were resuspended in PBS at a stock concentration of 1mM. Sequence and chemical modification of the lead candidate are provided below:

*DCLRE1C*-targeting ASO 1: [LR](A)[sP].[LR](T)[sP].[LR](A)[sP].[dR](G)[sP].[dR](T)[sP].[dR](T)[sP].[dR](G)[sP].[dR](G)[sP].[dR](A)[sP].[dR](T)[sP].[dR](A)[sP].[dR]([5meC])[sP].[dR](T)[sP].[LR]([5meC])[sP].[LR](G)[sP].[LR](G)

### Cell Culture

HAP1 WT and knockout cell lines (Horizon) were maintained in IMDM media supplemented with 10% heat-inactivated FBS and 1% penicillin/streptomycin. HEK293T (ATCC, CRL-3216) were maintained in DMEM-GlutaMAX with 10% fetal bovine serum (FBS) and 1% penicillin/streptomycin. HeLa cells (ATCC, CCL-2) were cultured in MEM-HEPES-GlutaMAX with 10% FBS and 1% penicillin/streptomycin. HepG2 (ATCC, HB-8065) were cultured in MEM-HEPES-GlutaMAX with 10% FBS, 1x non-essential amino acids, 1 mM sodium pyruvate and 1% penicillin/streptomycin. K562 cells (ATCC CRL-243) were maintained in RPMI-1640-GlutaMAX supplemented with 10% heat-inactivated FBS, 20 mM HEPES, 1 mM sodium pyruvate and 1% penicillin/streptomycin. Primary human hepatocytes (PHHs) from a deceased female donor (LifeNet Health, LOT: 2018274-01) were first thawed in pre-warmed Human Hepatocytes Thawing Medium (MED-HHTM4C-50ML, LifeNet Health), washed once with PBS, then re-suspended in 20 mL of Human Hepatocytes Plating Medium with supplement (MED-HHPM-250ML, MED-HHPMS-15ML, LifeNet Health). The following day, the medium was replaced with maintenance medium (William’s E medium, no phenol red (Gibco), 5% FBS, 0.1 µM Dexamethasone, 0.5% penicillin/streptomycin, 1X Insulin-Transferrin-Selenium (ITS, 41400-045 (Gibco), 2 mM L-Glutamine, and 15 mM HEPES pH 7.4) for the remainder of the experiment. hiPSCs (AstraZeneca-generated cell line, R-iPSC Clone J, LineID: iPS.1) were maintained in Cellartis DEF-CS Culture System (Cat. No. Y30010). hiPSCs-derived cardiomyocytes stored in cryovials were first thawed and seeded in customized thawing media (RPMI-1640 with B27 supplement and ascorbic acid) on fibronectin pre-coated plates. After three days of incubation, the cells were maintained in iCell Maintenance media (FUJIFILM) with frequent media changes. Cells were maintained at 37 °C in a 5% CO_2_ atmosphere and regularly tested for mycoplasma contamination. Cell line identity was authenticated through STR profiling (IDEXX BioAnalytics).

### KnockOut and KnockIn Cell Line Generation

*DCLRE1C* knockouts were generated in HEK293T and iPSCs with *Sp*Cas9 RNPs using a pooled three gRNA approach (Synthego, CRISPR Gene Knockout Kit from EditCo Bio). Spacer sequences used were: 1) 5’-UCACUCACCUUUGUGGCAGU-3’, 2) 5’-CUCCCUAUCGAAGCGGUCUA-3’, and 3) 5’-CGGCGCUAUGAGUUCUUUCG-3’. Briefly, RNP complexes were made by mixing equimolar *Sp*Cas9 protein (IDT, Alt-R™ *Sp*Cas9 Nuclease V3) and pooled gRNAs and electroporated into cells lines by Nucleofection (Lonza). For control, cell lines were also transfected with RNP complexes of *Sp*Cas9 and a non-targeting gRNA (spacer = 5’-GCGUCGUCGGUCGCGAUUAA-3’). For HEK293T, cells were electroporated with the Nucleofector 4D system (Lonza) using these conditions: 2E05 cells, RNP: *Sp*Cas9 (50pmol), gRNAs equimolar mix (50 pmol), electroporating enhancer oligo (50 pmol), CM-130 pulse program (Lonza). For iPSC, cells were electroporated with the Neon Nxt electroporation system (Thermo) using these conditions: 4.5E05 cells, RNP: *Sp*Cas9 (72pmol), gRNA equimolar mix (216 pmol), electroporating enhancer oligo (75 pmol), electroporation parameters: 1100V, 30ms, 1 pulse. After electroporation, cells were grown for three days before harvesting some cells for knockout validation, and expanding the rest of the cells to make freezer stocks. Knockout was confirmed by Sanger sequencing using Synthego ICE analysis tool (https://www.synthego.com/products/bioinformatics/crispr-analysis). Knockout pool generated for HEK293T and iPSCs had a calculated knockout score of 99% and 87%, respectively.

For overexpression of exogenous genes in HEK293T cell lines (Figure 1d, Figure 2d, and Supplementary Figure 2), cell lines were generated by Xential [13]. Template plasmids for integration into the *HBEGF* locus for Diphtheria Toxin (DT) selection was designed to contain a splicing acceptor sequence, followed by the mutated sequence of the *HBEGF* exon 4 linked to a self-cleaving peptide (T2A), followed by the doxycycline-inducible system all flanked by 750bp homology arms to the left and right of the *HBGEF* target site. The doxycycline-inducible system consists of the reverse tetracycline-controlled transactivator protein (rtTA) fused with a blasticidin resistance gene with a self-cleaving peptide (P2A), all under the expression of the endogenous *HBEGF* promoter, and then a downstream Tetracycline Response Element 3rd Generation (TRE3G) promoter to drive the expression of the human expression codon optimized sequence CDS of the gene of interest fused with GFP separated by a self-cleaving peptide (T2A). Sequences for repair templates can be found in Supplementary Data 3. All constructs were synthesized and cloned into a bacteria vector backbone by Genscript. HEK293T cells (WT or *DCLRE1C* knockout) were transfected with *Sp*Cas9 expressing plasmid, *HBEGF*-targeting gRNA (5’-GGGUGAUGUUGCCUGACCGG-3’), and repair templates with Fugene HD. Briefly, 800,000 cells were seeded into a 6-well plate in 2 mL of growth media one day prior to transfection. Then, 3000 µg of plasmid at a 1:1:2 ratio of *Sp*Cas9:gRNA:repair template was mixed with a Fugene HD reagent-to-DNA ratio of 0.6 in a total volume of 150 µL and incubated at room temperature for 15 min before adding to the cells. After 72 hours of incubation, DT selection was started. Cells were harvested from each 6-well plates and split 1/2 into a new 6-well plate with media containing 20 ng/µL of DT (The Native Antigen Company). After 48 hours, media was replaced with freshly prepared DT (20 ng/µL) and blasticidin (10 ng/µL) containing media. After which, freshly prepared DT- and blasticidin-containing media was replaced every two days, and cells harvested and split whenever wells became confluent, for 9 days in total and until control transfected cells were no longer proliferating or present. After selection, media was replaced twice with non-DT containing media on consecutive days, before expanding in a 10 cm dish and frozen into working stocks.

### HAP1 Knockout Screening and NHEJ Genes Overexpression Experiment

For the HAP1 knockout experiments, cells line vials ordered from Horizon Discovery (https://horizondiscovery.com/en/engineered-cell-lines/products/human-hap1-knockout-cell-lines) were thawed and expanded in culture dishes, maintaining <70% confluency (<2 passages) before freezing working stocks. Samples of each cells lines were tested for mycoplasma contamination, authenticated through STR profiling (IDEXX BioAnalytics), and knockouts validated by PCR using primers defined in the HAP1 cell line data sheets and comparing amplicon size with the WT background. For screen, working vials of all HAP1 cell lines were thawed and passaged twice in culture dishes maintaining <70% confluency before starting the experiment. One day prior to the experiment, 4 million cells of each cell lines were seeded on separate 10 cm dish. The next day, RNP complexes of PEn together with synthetic pegRNA, springRNA, or non-targeting gRNA were formulated by mixing, for each reaction, 50 pmols of protein together with 50 pmols or synthetic pegRNA or springRNA and 50 pmols of an electroporating enhancer oligo (5’-CCAGCAGAACACCCCCATCGGCGACGGCCCCGTGCTGCTGCCCGACAACCACTACCTGAGCACCCAGTCCGCCCTGAGCAAAGACCCCAACGAGA-3’) and incubated at room temperature for 10min before placing on ice. Cells were then harvested (TrypLE) and counted. Number of cells required for electroporation were collected by centrifugation (300xg, 5min), washed 1X with PBS (1mL), and resuspended in nucleofection buffer (SE Cell Line 96-well Nucleofector™ Kit; Lonza). For each reaction, 400,000 cells were resuspended in 16.4 µL of Nucleofector SE cell line solution and 3.6 µL Nucleofector Supplement. Subsequently, RNP complex (50:50:50 pmols/reaction or roughly 2 µL/reaction) were added to cells and gently mixed by pipetting. The final ∼22 µL suspensions were transferred into the electroporation cuvettes and electroporated with pulse code DS-118 (Lonza 4D-Nucleofector). Immediately after electroporation, 80 µL of pre-warmed media was added to the cuvettes to allow the cells to recover. Then 25 µL of cell mixture was added to a 96-well plate with 175 µL of pre-warmed media. Cells were collected for DNA extraction after 72 hours. This process was repeated twice for two biological replicates.

For the NHEJ genes overexpression experiment, 20,000 cells of each generated cell lines (HEK293T background) were seeded into wells of a 96-well plate in media containing 2 µg/mL of doxycycline to induce expression of NHEJ genes. The next day, the cells were transfected with FuGENE HD (Promega) using a 6:1 Fugene-to-DNA ratio and 100 ng of total DNA per well of 96-well plate (75 ng of editor and 25 ng of pegRNA/springRNA). After transfection, plates were incubated for 72 hours before harvesting the cells for DNA extraction. Additionally, RFP and GFP levels were monitored by fluorescent microscopy to confirm activation of the tetON promoter.

### DNA, RNA, Protein Delivery into Different Cell Types

This study employed multiple editor formats and guide modalities across adherent and suspension cell types. For consistency, we refer to the tethered nuclease-RT as PEn (RNP) or PEnMAX (plasmid), untethered format as Cas9 + MMLV RT, and to guide RNAs as pegRNAs (with homology arm) or springRNAs (without homology arm; PRINS). All sequences, chemistries, and translated protein sequences are provided in Supplementary Data 1–2. Delivery choices for each experiment are aligned to the figures noted below.

1) HAP1 cells

All experiments with HAP1 cells (Figure 1,2 Supplementary Figure 4) were completed by RNP electroporation of PEn protein complexed with synthetic pegRNA/springRNA and enhancer oligo, as described in the previous section. For HAP1 experiments with DNA-PKcs inhibitor, cells were directly seeded in AZD7648 (1 µM) or DMSO control containing media after electroporation and continued for 72 hours until harvest.

2) HEK293T cells

For experiments with HEK293T cells, different modalities and delivery methods were used. For plasmid only transfection (Figure 2d, Figure 3e,f, Supplementary Figure 2, Supplementary Figure 5a), 20,000 cells were seeded into wells of a 96-well plate in media. The next day, cells were transfected with FuGENE HD (Promega) using a 6:1 Fugene-to-DNA ratio and 100 ng of total DNA per well of 96-well plate (75 ng of editor and 25 ng of pegRNA/springRNA). After transfection, plates were incubated for 72 hours before harvesting the cells for DNA extraction. Where applicable (doxycycline-inducible overexpression), cells were seeded in media containing 2 µg/mL of doxycycline one day prior to plasmid transfection to induce expression of gene of interest.

For plasmid and RNA co-transfections in HEK293T (Figure 3 a,b,d, and Figure 5b), 12,500 cells were seeded into wells of a 96-well plate. The next day, cells were transfected with FuGENE HD (Promega) using a 6:1 Fugene-to-DNA ratio and 100 ng of total DNA per well of 96-well plate (100 ng of PEnMAX editor). One day post DNA transfection, cells were transfected with Lipofectamine RNAiMAX (Invitrogen) and 2 pmol of synthetic springRNAs. Briefly, 0.5 μL of RNAiMAX were mixed with 8.3 μL of Opti-MEM (Gibco) to form a working RNAiMAX solution. In parallel, 2 pmols of synthetic gRNA were mixed with 5 μL of Opti-MEM (Gibco) to form a working gRNA solution. 5 μL of working RNAiMAX solution were mixed with 5 μL of working gRNA solution, incubated at room temperature for 5 minutes and added to cells. After which, plates were incubated for 48 hours before harvesting the cells for DNA extraction.

For siRNA knockdown of different nucleases in HEK293T cells (Figure 3c), cells were transfected with 120 pmols of siRNA in 6-well plates by Lipofectamine RNAiMAX (Invitrogen) using 9 μL of transfection reagent per well. Then after three days, cells were harvested and re-seeded in a 96-well plate. The following day, cells were transfected with 200 ng of PEnMAX plasmid, 66 ng of springRNA plasmid, and 1 pmol of siRNA with Lipofectamine 2000 (Invitrogen) using 0.5 μL of transfection reagent per well. After which, plates were incubated for 72 hours before harvesting the cells for DNA extraction.

For initial CRISPRoff/siRNA/ASOs silencing/knockdown experiments in HEK293T cells (Figure 4a), all components were delivered by transfection. 500,000 HEK293T cells were reverse-transfected into wells of a 24-well plate. For epigenetic silencing, 500ng of *Sp*Cas9-CRISPRoff editor mRNA and 15 pmols of either non-targeting or *DCLRE1C-*targeting gRNA were delivered by Lipofectamine MessengerMax (Invitrogen) using 1.5 μL of transfection reagent per well. For siRNA and ASOs, 15 pmols of either non-targeting or *DCLRE1C-*targeting siRNA/ASO were delivered by Lipofectamine RNAiMAX (Invitrogen) using 1.5 μL of transfection reagent per well. Epigenetic silencing and plasmid delivery (Figure 4b), were completed by first transfecting 750,000 HEK293T cells (WT or *DCLRE1C* knockout) seeded into wells of a 6-well plate with the epigenome editor mRNA (2.5 µg/ well) and either non-targeting or *DCLRE1C* targeting gRNA (50pmols/ well) by Lipofectamine MessengerMax (7.5μL/ well). After five days of incubation, the cells were re-seeded into a 96-well plate (12,500 cells /well). The following day, cells were transfected with FuGENE HD (Promega) using a 6:1 Fugene-to-DNA ratio and 100 ng of total DNA per well of 96-well plate (75 ng of editor and 25 ng of springRNA). After transfection, plates were incubated for 72 hours before harvesting the cells for DNA extraction.

3) HEK293T, HeLa, HepG2, and K562 (CRISPRoff silencing and ASOs knockdown)

For subsequent epigenetic silencing experiments (Figure 4c, Supplementary Figure 8), target cells were electroporated with the CRISPRoff editor mRNA and either non-targeting or *DCLRE1C-*targeting gRNA to improve silencing efficiency. HEK293T, HeLa, HepG2, and K562 cells were collected, washed once with PBS, and resuspended in nucleofection buffer (SF Cell Line 96-well Nucleofector™ Kit; Lonza) containing 2 μg of *Sp*Cas9 epigenome editor mRNA and 100 pmols of either the non-targeting or *DCLRE1C*-targeting gRNA (2E05, 2E05, 3E05, and 3E05 cells/condition, respectively). Cell mix were electroporated using the Nucleofector 4D unit (Lonza) and the CM-130 (HEK293T), CN-114 (HeLa), EH-100 (HepG2) or FF-120 (K562) pulse code. Post-electroporation, cells were transferred into 2 mL of pre-warmed medium in 6-well plates and incubated for at least 5 days before being passaged in fresh media and grown to confluency before continuing with the electroporation of the PRINS components. Cells were once again collected, washed once with PBS, and resuspended in nucleofection buffer (SF Cell Line 96-well Nucleofector™ Kit; Lonza) containing 2 μg of PEn mRNA, 1 μg of La(1-194) mRNA, and 100 pmols of each springRNA (2E05, 2E05, 3E05, and 3E05 cells/condition, respectively). Cell mix were electroporated using the Nucleofector 4D unit (Lonza) and the CM-130 (HEK293T), CN-114 (HeLa), EH-100 (HepG2) or FF-120 (K562) pulse code. Post-electroporation, cells transferred into 200 µL of pre-warmed media in 96-well plates and incubated for 72 hours before harvesting the cells for DNA extraction.

For ASOs knockdown experiments in different cell types (Figure 4d, Figure 5c,d,e, Supplementary Figure 9), target cells were electroporated with either the non-targeting or *DCLRE1C-*targeting ASOs. To initiate knockdown, HEK293T, HeLa, and K562 cells were collected, washed once with PBS, and resuspended in nucleofection buffer (SF Cell Line 96-well Nucleofector™ Kit; Lonza) containing 100 pmols of either the non-targeting or *DCLRE1C*-targeting ASOs (2E05, 2E05, and 3E05 cells/condition, respectively). Cell mix were electroporated using the Nucleofector 4D unit (Lonza) and the CM-130 (HEK293T), CN-114 (HeLa), CN-114 (HeLa), or FF-120 (K562) pulse code. Post-electroporation, cells were transferred into 2mL of pre-warmed medium in 6-well plates and incubated for three days before continuing with the electroporation of the PRINS components. Cells were once again collected, washed once with PBS, and resuspended in nucleofection buffer (SF Cell Line 96-well Nucleofector™ Kit; Lonza) containing 2 μg of PEn mRNA, 1 μg of La(1-194) mRNA, and 100 pmols of each springRNA (2E05, 2E05, and 3E05 cells/condition, respectively). Cell mix were electroporated using the Nucleofector 4D unit (Lonza) and the CM-130 (HEK293T), CN-114 (HeLa), or FF-120 (K562) pulse code. Post-electroporation, cells dedicated for DNA extraction or HIBIT assay were transferred (1/5 of total cells) into 200 µL of pre-warmed media in 96-well plates and incubated for 72 hours before harvesting the cells or conducting the HIBIT lytic assay. Cells dedicated for protein extraction were instead transferred into first 24-well plates, then expanded into 6-well plates until 2 wells of a plate were confluent (1-2 weeks) before harvesting the cells.

4) Primary Human Hepatocytes

For work in primary human hepatocytes (Figure 5b), cells were thawed and seeded at 20,000 cells per well of a 96-well plate. After one day incubation, the media was replaced with fresh media (as described in the‘Cell Culture’ section. Primary cells were transfected with 168 ng of PEnMAX mRNA and 10 pmols of synthetic springRNA per well with Lipofectamine MessengerMax (Invitrogen) using 0.3 µL of transfection reagent per well. Cells were incubated for three days before measuring HIBIT activity.

5) iPSCs and iPSC-derived cardiomyocytes

For iPSC WT vs *DCLRE1C* knockout experiment (Supplementary Figure 5b), cells were electroporated with RNPs of PEn protein complexed with synthetic springRNA and oligo enhancer. Briefly, 450,000 cells were mixed with pre-complexed RNPs consisting of 72 pmols of PEn protein, 216 pmols of springRNA, and 75 pmols of oligo enhancer, then electroporated with the Neon Nxt electroporation system (Thermo) using these parameters: 1100V, 30ms, 1 pulse. Then cells were seeded into pre-coated plates and incubated for 72 hours before harvesting for DNA extraction.

For hiPSC-derived cardiomyocytes (Figure 5f), cells were first treated with 1.5 pmols of either non-targeting or *DCLRE1C-*targeting ASO with Lipofectamine RNAiMAX (Invitrogen) using 0.3 μL of transfection reagent per well. After three days of incubation, some cells were collected for RNA extraction and the remaining cells were transfected with 75ng of PEnMAX mRNA and 1.5 pmols of synthetic springRNA with Lipofectamine MessengerMax (Invitrogen) using weight RNA/volume transfection reagent ratio of 0.67. Cells were incubated for three days before measuring HIBIT activity.

### Genomic DNA Extraction and Amplicon Sequencing

Cells were harvested using Quick Extract solution (Lucigen) or Fast Extract solution (VWR) by directly adding 50 µL of DNA extraction solution per well of a 96-well culture plate with media removed, incubated for 10min at 37°C, then scratched off and transferred into a 96-well PCR plate. For K562 cells (suspension cells), cells were first centrifuged for 5 min at 300xg before removing the media and adding the DNA extraction solution. PCR plates were sealed, vortexed (10 sec), centrifuged, before adding to the thermocycler: 70°C for 10 min, 98°C for 10 min. Extracted DNA was stored in the freezer (-20°C) until needed.

Amplicons were generated with Phusion Flash High-Fidelity 2x Mastermix (F548, Thermo Scientific) or Q5 Hot Start High-Fidelity 2x Mastermix (M0492, NEB) in a 15 μL reaction, containing 1.5-2 μL of genomic DNA extract and 0.2 μM of target-specific primers with barcodes and adapters for next generation sequencing (NGS). All primer sequences are listed in Supplementary Data 4. PCR cycling conditions for Phusion Flash High-Fidelity 2x Mastermix were: 98 °C for 3 minutes, followed by 32 cycles of 98 °C for 10 seconds, 60 °C for 5 seconds, and 72 °C for 5 seconds. For Q5 Hot Start High-Fidelity 2x Mastermix the following PCR protocol was applied: 98 °C for 3 minutes, followed by 30 cycles of 98 °C for 10 seconds, 60 °C for 20 seconds, and 72 °C for 30 seconds, and final elongation at 72 °C for 2 minutes. All amplicons were purified using HighPrep PCR Clean-up System (MagBio Genomics). The size, purity, and concentration of amplicons were determined using a fragment analyzer (Agilent). To add Illumina indexes to the amplicons, samples were subjected to a second round of PCR. Indexing PCR was performed using KAPA HiFi HotStart Ready Mix (Roche), 0.067 ng of PCR template and 0.5 µM of indexed primers in the total reaction volume of 25 µL. PCR cycling conditions were 72 °C for 3 minutes, 98 °C for 30 seconds, followed by 10 cycles of 98 °C for 10 seconds, 63 °C for 30 seconds, and 72 °C for 3 minutes, with a final extension at 72 °C for 5 minutes. Samples were purified with the HighPrep PCR Clean-up System (MagBio Genomics) and analyzed using a fragment analyzer (Agilent). Samples were quantified using a Qubit 4 Fluorometer (Life Technologies) and subjected to sequencing using Illumina NextSeq system according to the manufacturer’s instructions.

### Bioinformatic Analysis

Demultiplexing of the amplicon-seq data was performed using bcl2fastq software. The fastq files were analyzed using CRISPResso2 V2.2.12. Detailed parameters are listed in Supplementary Data 5. Indel histograms and intended or longer insertion frequencies were analyzed and generated from the CRISPResso2 analysis files using a custom R-script and ggplot2. For indel histograms,‘Reference’,‘Prime-edited’, and‘Scaffold-incorporated’-aligned indel histograms from each samples were concatenated together adjusting the indel frame of reference for the‘Prime-edited’ and‘Scaffold-incorporated’ by the size of the springRNA insert (e.g. indel size = 0-> indel size = springRNA insert size) between indel sizes of-200 to + 200. Reads per indel size were normalized to the total number of reads per sample and multiplied by 100%. Indel histogram plots were generated using ggplot2 package, taking the average of the biological replicates for each indel size. Intended and longer insertion frequencies were calculated using the indel histograms, plotting the average of the biological replicates of the intended insertion length or the sum of all longer-than-intended insertions for each condition.

### RNA extraction and qPCR

RNA from the cells harvested from knockdown experiments was extracted using RNeasy Mini Kit (Qiagen) and quantified with a spectrophotometer. Then, 1000ng of RNA was converted into cDNA using the High-Capacity cDNA Reverse Transcription Kit (Applied Biosystems) in 20µL reactions. For qPCR, *DCLRE1C* expression was quantified by pre-designed primer/probe Taqman assay (Assay ID: Hs01052788_m1, Gene Symbol: *DCLRE1C*, Dye Label and Assay Concentration: FAM-MGB / 60X, Applied Biosystems) and normalized to *TBP* reference gene Taqman assay (Assay ID: Hs99999910_m1, Gene Symbol: *TBP*, Dye Label and Assay Concentration: FAM-MGB / 20X, Applied Biosystems). Briefly, 3µL of cDNA samples was added to a 10 µL reaction with TaqMan Fast Advanced Master Mix (Applied Biosystems) and appropriately diluted (1X) Taqman assay reagent. *DCLRE1C* and *TBP* assays were run separately, and all samples were run in technical duplicates. The plates were cycled in the QuantStudio 7 Flex system (Applied Biosystems) and the qPCR cycling conditions for TaqMan Fast Advanced Mastermix were: 95 °C for 20 seconds, followed by 40 cycles of 95 °C for 1 seconds and 60 °C for 20 seconds. Analysis was done by using the delta delta Ct equation.

### HIBIT detection assay

HIBIT was detected using the Nano-Glo HIBIT Lytic Detection System (Promega) following the manufacturer’s instructions. Briefly, media was removed and replaced with 100 µL of PBS. For K562 cells, an additional centrifugation step to pellet the cells (500xg, 3min) was performed before removing the media. Then, 100 µL of Nano-Glo HIBIT Lytic Detection Reagent was added cells and by pipetting several times. After 10 min incubation, cell mixture was transferred to a black-well plate (if not already in one) and luminescence was recorded on a PheraStar FSX plate reader (BMG Labtech) with 1 second integration time.

### Protein extraction and Western Blotting

Cells harvested for Western blotting were lysed in RIPA buffer with additional proteinase and phosphatase inhibitors (Thermo). Combined pellets collected from two confluent wells of a 6-well plate, 120 µL of the lysis buffer was added to the pellets and vortexed for 30 seconds before incubating on ice for 10 minutes. Then, the cells were sonicated three times for five seconds at 20% power. Protein was quantified using the Pierce BCA Protein Assay Kit (Thermo). Proteins were diluted between 1 and 2 µg /µL in NuPAGE LDS sample buffer with in NuPAGE sample reducing agent then incubated for 10 minutes at 70 °C. Samples were then loaded on NuPAGE™ Bis-Tris Mini Protein Gels (12%) (Thermo) with Precision Plus Protein Dual Color Standards (Biorad). Each individual Western blot images in Figure 5 were run on separate gels. For gels to measure Gapdh and Beta actin protein levels, 20 µg of proteins were loaded per lane while for gels to measure MIF protein and the protein tags (HIBIT, FLAG, Myc, 6XHIS), 30 µg of proteins were loaded per lane. Gels were run by electrophoresis with NuPAGE MES SDS Running Buffer with additional NuPAGE Antioxidant, 120V for 1 hour. After electrophoresis, gels were transferred using the iBlot 2 Dry Blotting System on nitrocellulose membranes (7 min, 15V). After transfer, membranes were blocked for 30 minutes in 5% milk in TBS-T. Primary antibody incubation was left overnight at 4 °C on a plate rocker and secondary antibody was left for 1 hour at room temperature on a plate rocker. Primary antibodies used (and concentrations): rabbit anti-Gapdh mAb (1/1000, Cell signaling #5174) or rabbit anti-Gapdh pAb (1/1000, Thermo PA1987), rabbit anti-Beta Actin mAb (1/1000, Cell Signaling #4970), rabbit anti-MIF pAb (1/1000, Thermo PA5-27343), rabbit anti-FLAG mAb (1/1000, Cell Signaling #14793), rabbit anti-Myc tag mAb (1/1000, Cell Signaling, #2278), and rabbit anti-His tag mAb (1/1000, Cell Signaling, #12698). Secondary antibody used: HRP-conjugated Goat anti-Rabbit IgG (H+L) (1/2000, Thermo 31460). HRP detection was done with SuperSignal™ West Femto Maximum Sensitivity Substrate (Thermo) and ChemiDoc imaging system (Biorad). Brightness and contrast were adjusted on the raw images using ImageJ.

### Statistics & reproducibility

Data visualization and statistical analysis were conducted using GraphPad Prism 9 (GraphPad Software, Inc.). Figure legends contain information on statistical tests, sample sizes, and P-values. No data were excluded from the analyses. No statistical method was used to predetermine sample size. The experiments were not randomized. The investigators were not blinded during experiments and outcome assessment.

## Acknowledgements

We thank the AstraZeneca Discovery Sciences Genome Engineering team for support and input on this work, in particular Isabel Weisheit for development and testing of in-house mRNA epigenetic silencer. We thank Promega Corporation for sponsoring this project (L.C.D. is funded by Promega Corporation, M.S. and T.M. are employees of Promega Corporation). We thank Shalini Andersson and Steve Rees for supporting this project. We are grateful to all internal AstraZeneca teams for their contributions on NGS amplicon sequencing and analysis, protein synthesis, mRNA synthesis, and ASOs design and synthesis.

## Author Contributions

L.C.D. and M.M. conceptualized the study and prepared the manuscript. L.C.D., OK.C., A.M., A.L., and N.A. performed most of the experimental work and analysis with support from J.K., P-P. H., M.P., S.Sv, and G.S.. As well, NGS short amplicon sequencing was performed in large part by A.B. and J.L. as well as other internal team members. Sequencing data analysis pipeline and QC validation was done by M.F.. PEn protein synthesis and purification was done by E.G. mRNA design and synthesis was done by S.M. and G.T.. ASOs design was done by D.C. and F.K., and ASOs synthesis was done by S.Sa. and A.D.

---

**Supplementary Figure 1.**
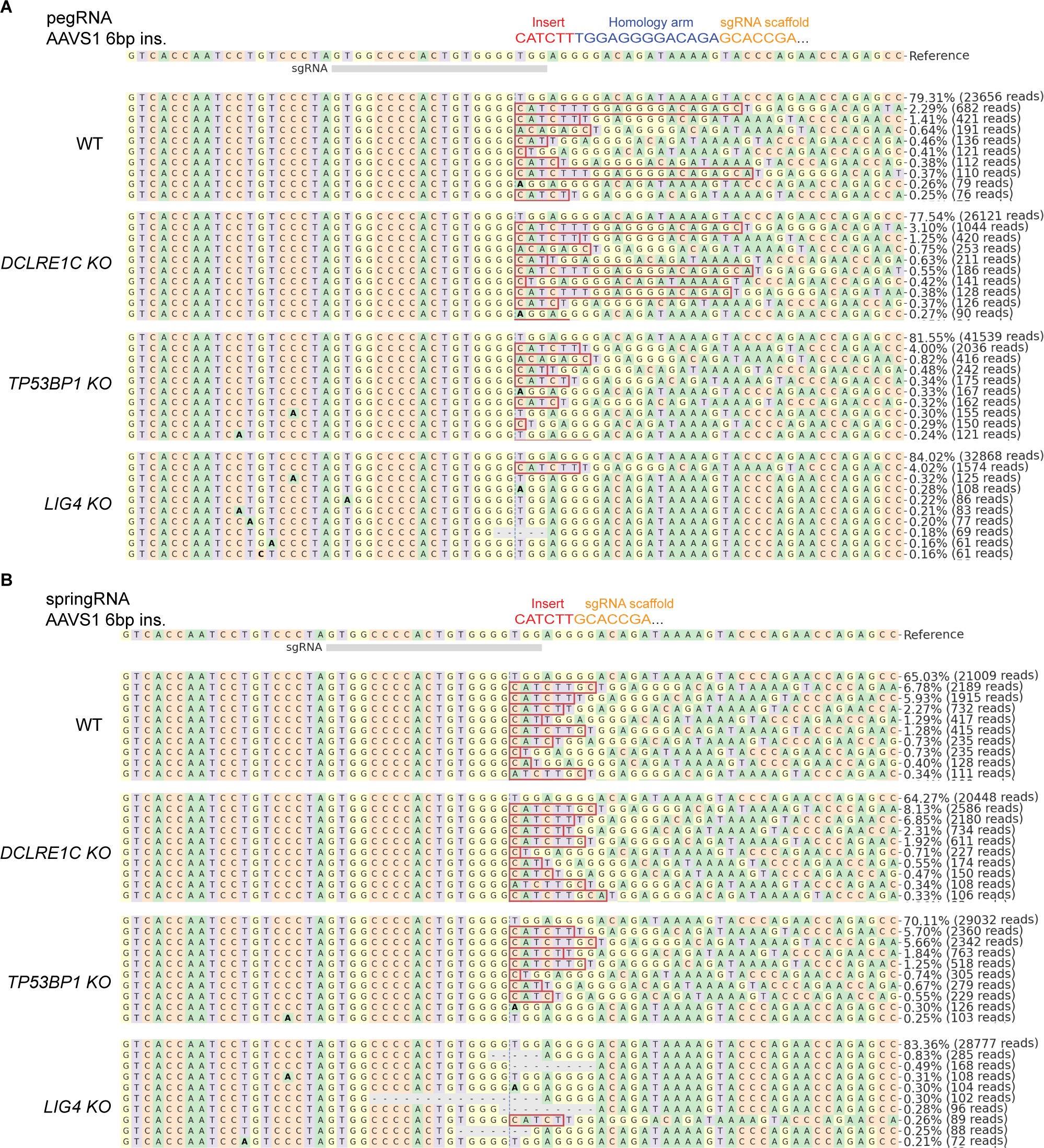
Allele Frequency Plots of HAP1 Knockout Targeted Screening with PEn. Representative allele frequency plots from experiment described in Figure 1a,b of the top 10 sequenced alleles between WT HAP1 and *DCLRE1C, TP53BP1,* and *LIG4* knockouts after editing with PEn and with A) pegRNA or B) springRNA into AAVS1 (*PPP1R12C*) target site with an intended insert of 6bp. Allele frequency plots were generated by CRISPResso2.

**Supplementary Figure 2.**
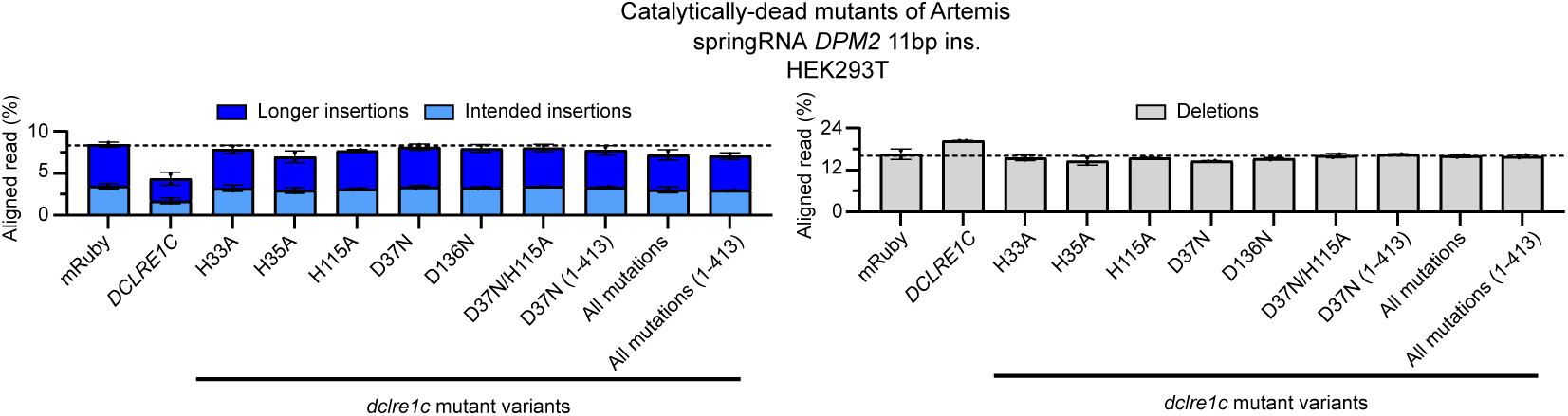
Effect of Overexpressing Catalytically Dead Variants of Artemis on PRINS Editing Outcomes. WT HEK293T cell lines were generated by Xential (see methods) with a doxycycline-inducible system to overexpress either exogenous mRuby (control) or *DCLRE1C* gene, or different variants catalytically dead variants of Artemis (*dclre1c* mutant variants). Cells were first treated with doxycycline (2 µg/mL) for one day to induce expression of cDNA, then transfected by lipofection with *Sp*Cas9-RT (PEnMAX) expressing plasmid and springRNA expressing plasmid for *DPM2* target site (11bp intended insertion). Bar graph shows average frequency (n=3 biological replicates ± SD) of intended or longer insertions (left) and deletions (right) calculated using by CRISPResso2 analysis and a custom R-script.

**Supplementary Figure 3.**
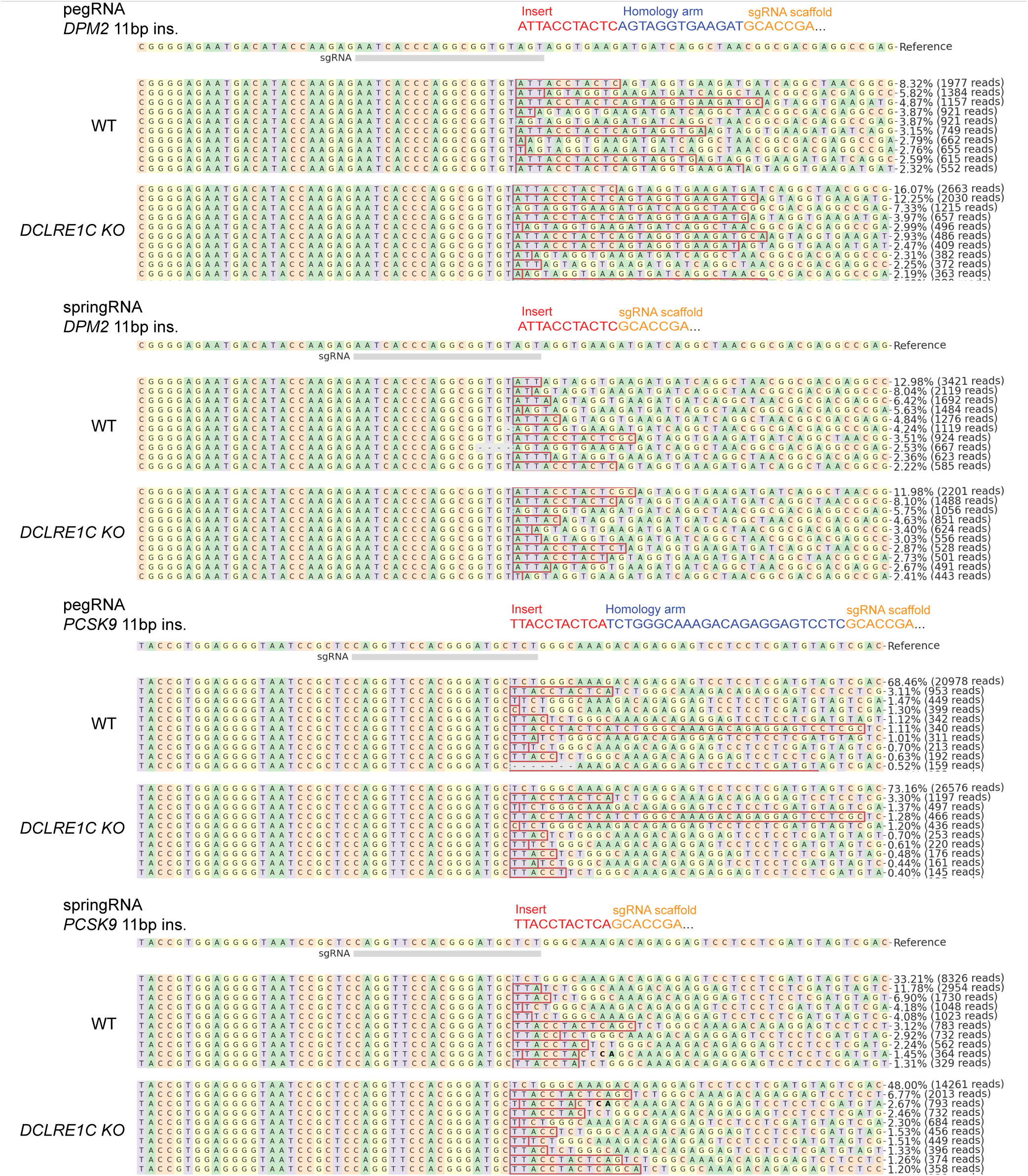
Allele Frequency Plots of WT and *DCLRE1C* Knockout HAP1 Cells Edited with PRINS for Longer Insert Sizes. Representative allele frequency plots from experiment described in Figure 2a,b of the top 10 sequenced alleles between WT and *DCLRE1C* knockout HAP1 cells after editing with PEn and with pegRNA or springRNA designed for insertion into *DPM2* (11bp insertion) or *PCSK9* (11bp insertion) target sites. Allele frequency plots were generated by CRISPResso2.

**Supplementary Figure 4.**
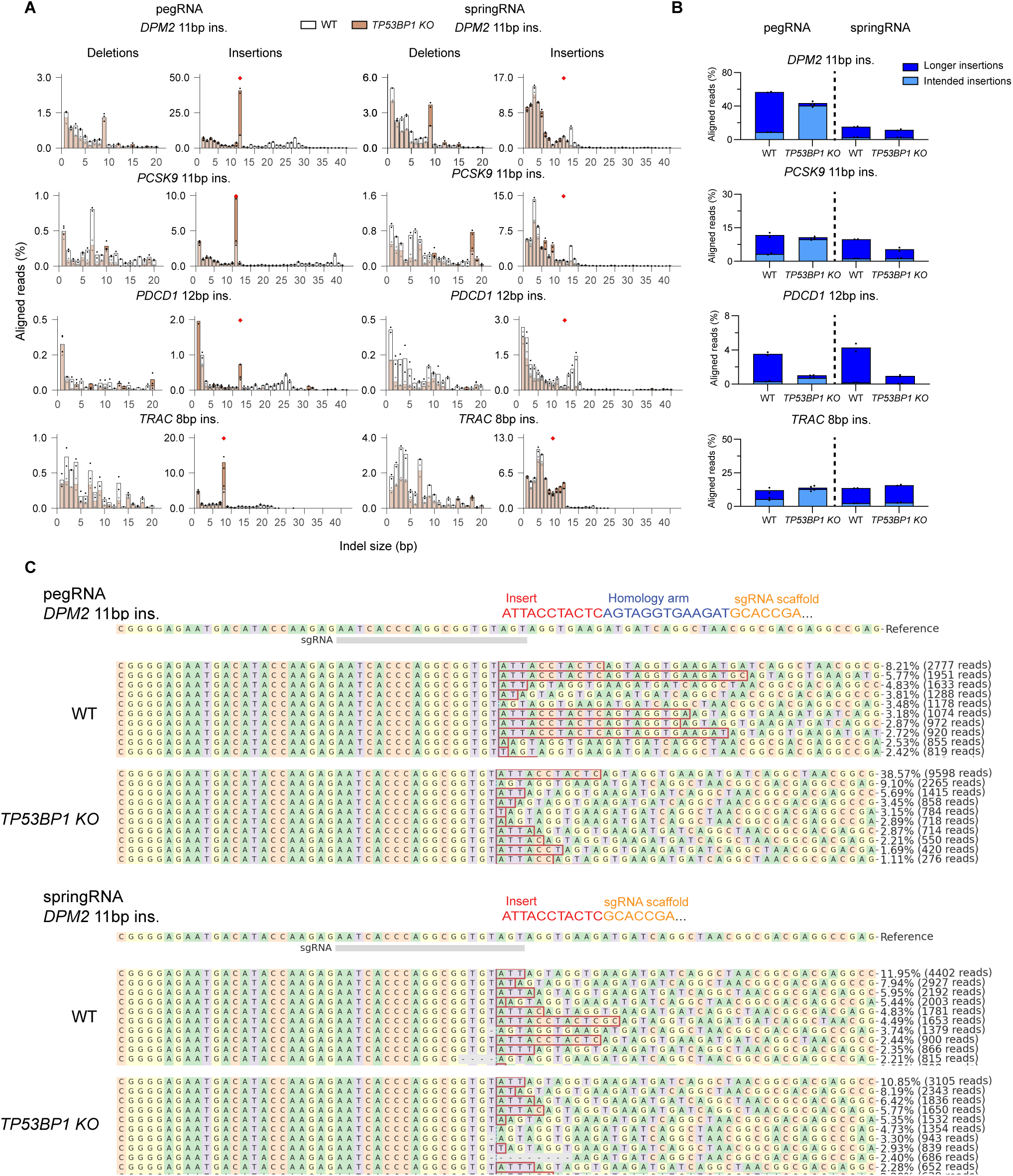
*TP53BP1* Knockout Increases the Precision of PEn-mediated pegRNA Insertions. A) Indel distribution patterns for PEn editing (with pegRNA or springRNA) in WT and *TP53BP*1 knockout HAP1 cells. Cells were electroporated with *Sp*Cas9-RT (PEn) RNPs and either pegRNA or springRNA for four target sites (*DPM2*-11bp, *PCSK9*-11bp, *PDCD1*-12bp, *TRAC*-8bp). Overlayed indel histograms (CRISPResso2, custom R-script) show average indel size frequencies (two biological replicates) comparing WT (white) and *TP53BP1* knockout (orange). Deletions (1-20 bp) are plotted on the left, and insertions (1-40 bp) on the right. Intended insertion size is indicated by a red diamond. B) Insertion frequencies of PEn editing between WT and *TP53BP1* knockout HAP1 cells. Bar graph shows average frequency (two biological replicates ± SD) of intended or longer insertions from the experiment described in (A), calculated using CRISPResso2 and a custom R-script. C) Representative allele frequency plots from experiment described above of the top 10 sequenced alleles between WT and *TP53BP1* knockout HAP1 cells after editing with PEn and with pegRNA or springRNA designed for insertion into *DPM2* (11bp insertion) target site. Allele frequency plots were generated by CRISPResso2.

**Supplementary Figure 5.**
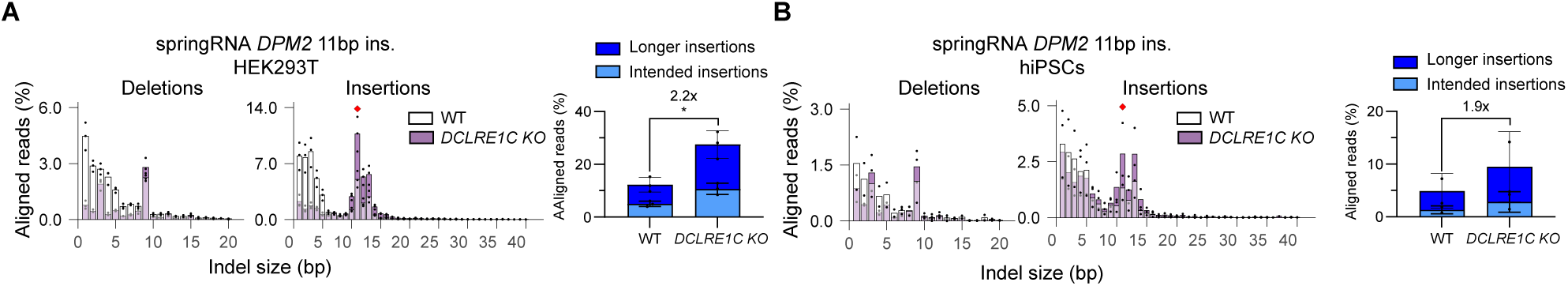
*DCLRE1C* Knockout Increases PRINS-mediated springRNA Insertions in HEK293T and iPSCs. Indel distribution patterns for PRINS editing with springRNA in A) WT HEK293T and B) iPSC cells, comparing to *DCLRE1C* knockout cells. For HEK293T cells, PEnMAX plasmid and springRNA plasmid (11bp insertion) at the *DPM2* target site were introduced via transfection. For hiPSC cells, PEn RNPs and springRNA (11bp *DPM2* insertion) were delivered via electroporation. Overlayed indel histograms (CRISPResso2, custom R-script) show average indel size frequencies for HEK293T (three biological replicates) and iPSCs (two biological replicates) comparing WT (white) and *DCLRE1C* knockout (purple) backgrounds. Intended insertion size is indicated by a red diamond. Bar graph shows average frequency (n=3 biological replicates ± SD) of intended or longer insertions calculated using CRISPResso2 and a custom R-script. *P*-values from Supplementary Figure 5 were determined using t-test (two-tailed, unpaired): * p < 0.05, ** p < 0.01, *** p < 0.001, **** p < 0.0001.

**Supplementary Figure 6.**
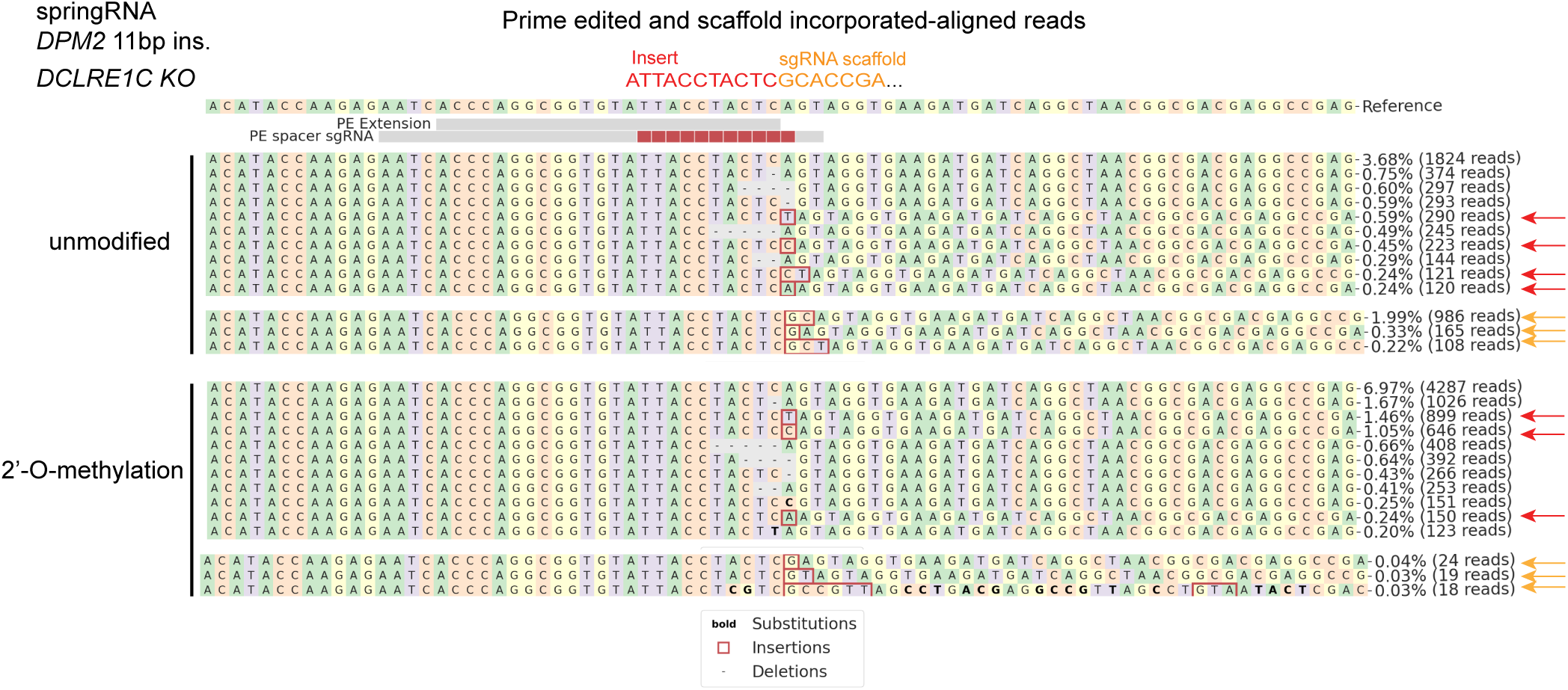
Allele Frequency Plots of *DCLRE1C* Knockout HEK293T Cells Edited with PRINS and Unmodified or 2’-O-methylation Modified springRNA. Representative allele frequency plots from experiment described in Figure 3d of the prime-edited aligned or scaffold-incorporated aligned alleles in *DCLRE1C* knockout HEK293T cells after editing with PEnMAX and unmodified or 2’-O-methylation modified springRNA designed for insertion into *DPM2* (11bp insertion) target sites. Allele frequency plots were generated by CRISPResso2. Red arrows indicated non-templated insertions and orange arrows indicated gRNA scaffold templated longer insertions into target site after the intended insertion.

**Supplementary Figure 7.**
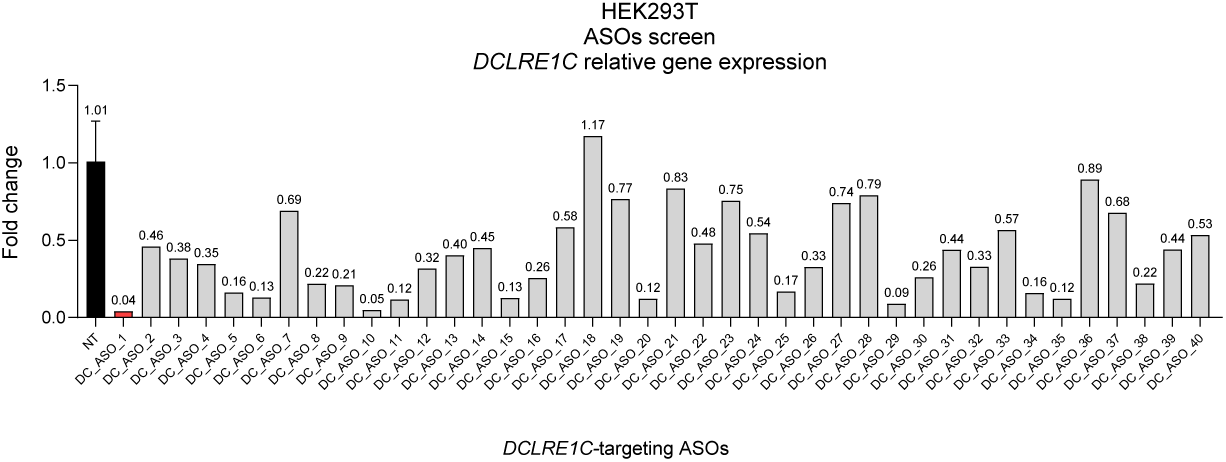
Screening Antisense Oligonucleotides (ASOs) for Targeted Knockdown of Artemis *(DCLRE1C*). HEK293T cells were subjected to ASO-mediated knockdown of different oligonucleotides targeting Artemis (*DCLRE1C*), alongside non-targeting (NT) ASO control. Relative fold change within cell line was calculated by delta delta Ct method using the non-targeting ASO as the control group and *TBP* as the reference gene. Bar graph show fold change for one biological replicate. Red bar (DC_ASO_1) was the lead candidate taken forward for all subsequent ASOs-mediated knockdown.

**Supplementary Figure 8.**
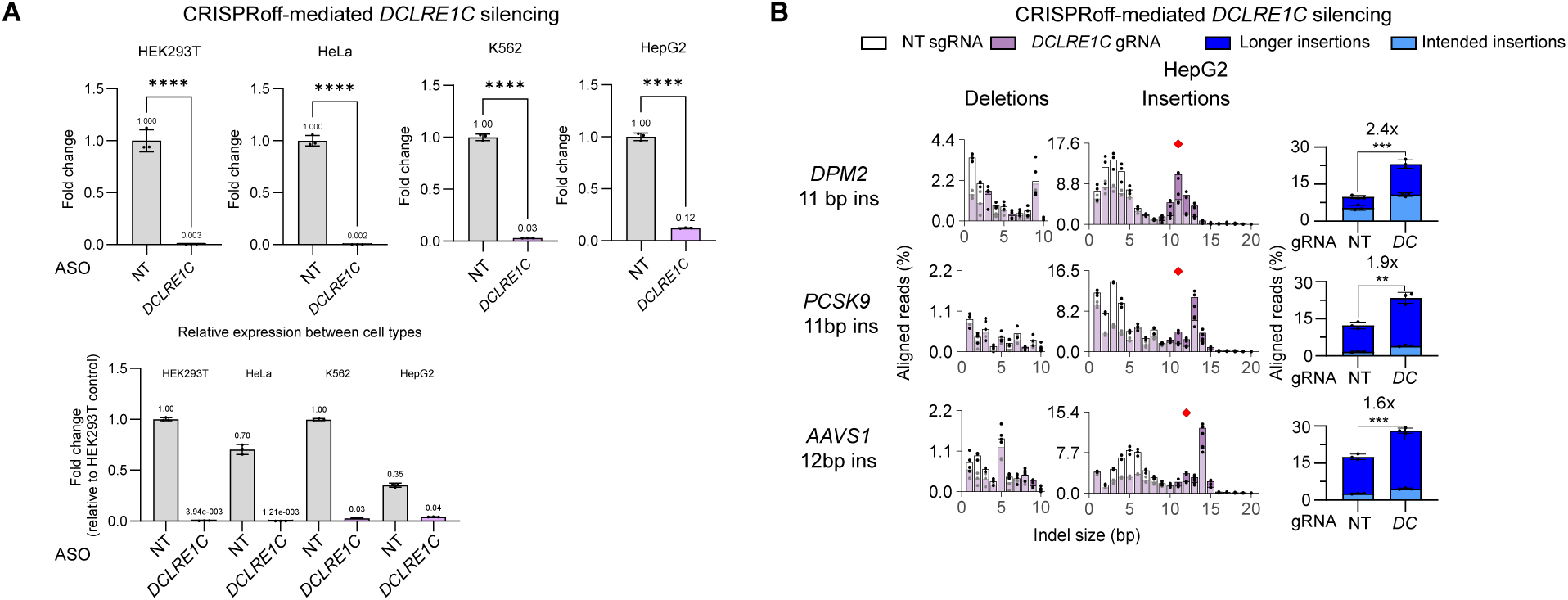
Epigenetic Silencing of Artemis (*DCLRE1C*) in Different Cell Lines and Effect on PRINS Editing in HepG2. A) Relative expression levels of Artemis gene (*DCLRE1C*) following CRISPRoff-mediated silencing prior to delivery of the PRINS components. RNA was extracted and reverse-transcribed into cDNA, before qPCR analysis using primers and probes specific for *DCLRE1C* and *TBP* (reference gene). Relative fold change within cell line was calculated by delta delta Ct method using the non-targeting silencing gRNA as the control group and *TBP* as the reference gene. Relative expression levels between all cell lines were calculated by delta Ct method using the non-targeting silencing gRNA in HEK293T cells as the control group. Bar graphs show average fold change for three biological replicates (n=3 biological replicates ± SD). B) CRISPRoff-mediated Artemis gene (*DCLRE1C*) silencing for PRINS insertion in HepG2 cells. Hepg2 cells were treated with CRISPRoff components (*DCLRE1C*-targeting or non-targeting gRNA), then transfected with PEnMAX mRNA, La(1-194) mRNA, and synthetic springRNAs for *DPM2*, *PCSK9*, and AAVS1 (*PPP1R12C*) insertions. Indel histograms (CRISPResso2, custom R-script) show average indel frequencies (n=3 biological replicates ± SD) for non-targeting (white) and *DCLRE1C*-targeting gRNA (purple) conditions for HepG2 cell background. Intended insertion size is indicated by a red diamond. Bar graph shows average frequency (n=3 biological replicates ± SD) of intended or longer insertions for all targets calculated using the analysis described above and a custom R-script. *P*-values from Supplementary Figure 8 were determined using t-test (two-tailed, unpaired): * p < 0.05, ** p < 0.01, *** p < 0.001, **** p < 0.0001.

**Supplementary Figure 9.**
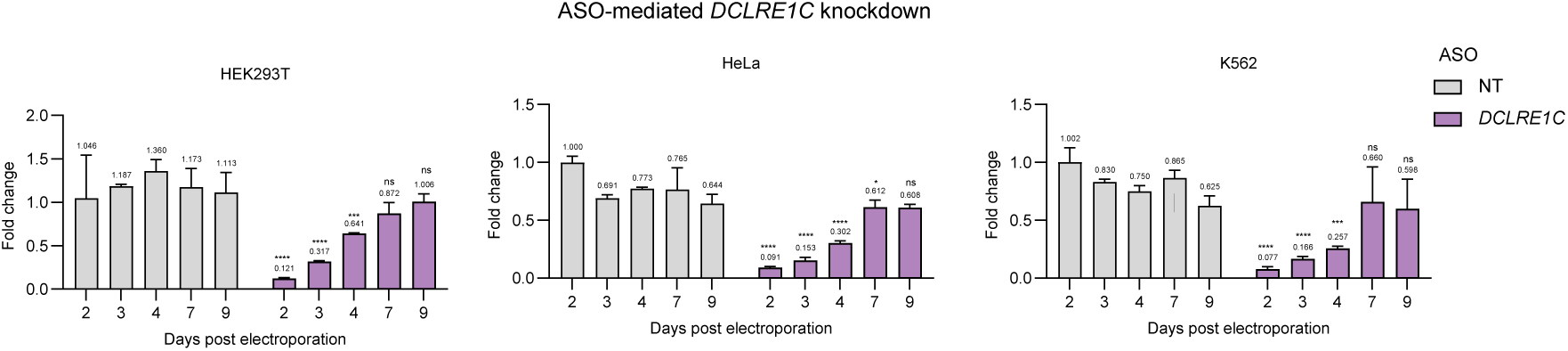
Time course for ASOs-mediated Knockdown in Different Cell Lines. HEK293T, HeLa, and K562 cells were electroporated with *DCLRE1C*-targeting ASO, alongside non-targeting (NT) ASO control. RNA was collected and *DCLRE1C* expression was measured by qPCR at indicated time points (Day 2, 3, 4, 7, 9) using delta delta Ct method using the non-targeting ASO as the control group and *TBP* as the reference gene. Bar graph shows average fold expression change of *DCLRE1C* relative to Day 2 non-targeting conditions for each cell line (n=3 biological replicates ± SD). *P*-values from Figure 4b were determined using ANOVA two-way test with multiple comparisons using Tukey’s test. Displayed pairwise comparisons are relative to NT of the equivalent time point: * p < 0.05, ** p < 0.01, *** p < 0.001, **** p < 0.0001.

